# Ubiquitous selfish toxin-antidote elements in *Caenorhabditis* species

**DOI:** 10.1101/2020.08.06.240564

**Authors:** Eyal Ben-David, Pinelopi Pliota, Sonya A. Widen, Alevtina Koreshova, Tzitziki Lemus-Vergara, Philipp Verpukhovskiy, Sridhar Mandali, Christian Braendle, Alejandro Burga, Leonid Kruglyak

**Affiliations:** Department of Human Genetics, Department of Biological Chemistry, and Howard Hughes Medical Institute, University of California, Los Angeles, CA 90095, USA; Department of Biochemistry and Molecular Biology, Institute for Medical Research Israel-Canada, The Hebrew University School of Medicine, Jerusalem, Israel; Institute of Molecular Biotechnology of the Austrian Academy of Sciences (IMBA), Vienna BioCenter (VBC), Dr. Bohr-Gasse 3, 1030 Vienna, Austria; Department of Biological Chemistry, University of California, Los Angeles, CA 90095, USA; Université Côte d’Azur, CNRS, Inserm, IBV, Nice 06100, France

## Abstract

Toxin-antidote elements (TAs) are selfish genetic dyads that spread in populations by selectively killing non-carriers. TAs are common in prokaryotes, but few examples are known in animals. We discovered five maternal-effect TAs in the nematode *Caenorhabditis tropicalis* and one in *C. briggsae*. Unlike previously reported TAs, five of these novel toxins do not kill embryos but instead cause larval arrest or developmental delay. We identified the genes underlying a TA causing developmental delay, *slow-1/grow-1*, and found that the toxin, *slow-1,* is homologous to nuclear hormone receptors. Last, we found that balancing selection of conflicting TAs hampers their ability to drive in populations, leading to more stable genetic incompatibilities. Our results show that TAs are common in *Caenorhabditis* species, target a wide range of developmental processes, and may act as barriers preventing gene flow.

## Introduction

Selfish genetic elements promote their own survival at the expense of their host’s fitness (*1*–*3*). As a result of this conflict, selfish elements underlie numerous cases of hybrid dysgenesis, sterility, and genetic incompatibility in the wild (*4*–*7*). Toxin-antidote elements (TAs) are an extreme example of selfish elements—they spread in natural populations by killing non-carrier individuals. TAs comprise two tightly linked genes: a toxin and its cognate antidote (Fig. 1A). The toxin is expressed in one of the two gametes, whereas the antidote is expressed zygotically. In crosses between individuals that carry the TA and ones that do not, the toxin is delivered to all of the progeny by the egg or sperm, and only embryos that inherit at least one copy of the TA survive because they also express the antidote (Fig. 1A).

**Figure 1.**
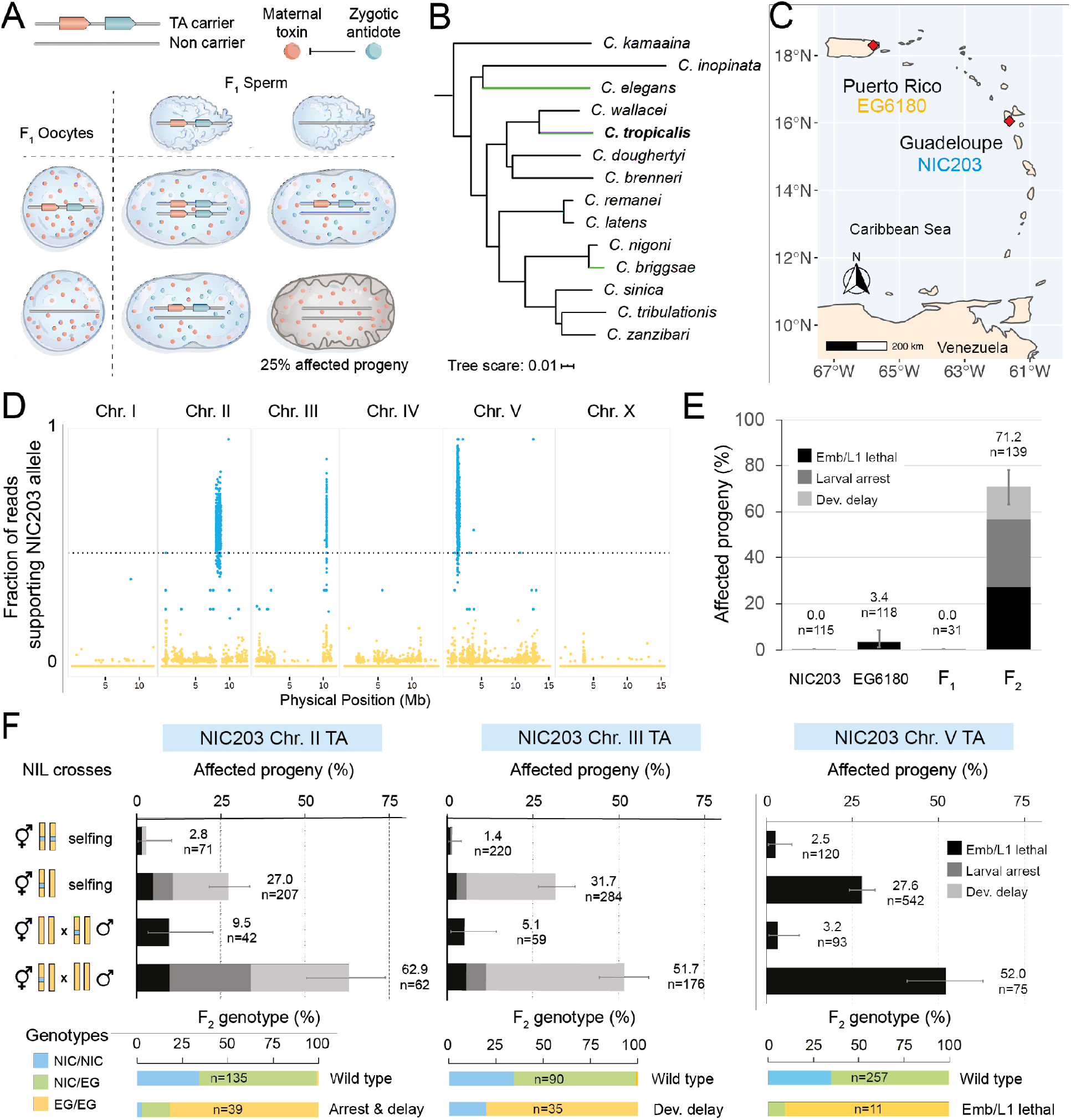
Multiple maternal-effect TAs are active in *C. tropicalis* and target embryonic and post-embryonic development. **(A)** A model for a maternal-effect Toxin-Antidote element (TA). The toxin (salmon circle) is expressed in the germline and deposited into eggs, whereas its antidote (cyan circle) is only expressed zygotically. Punnett square depicts the expected percentage of affected F_2_ progeny for a fully penetrant TA. **(B)** The phylogenetic relationship of the nematodes in the Elegans group. The phylogenetic tree has been adapted from the work of Stevens et al. (2019) (*18*). Hermaphroditic species, including *C. tropicalis* (bold), are highlighted with green branches. Scale is substitutions per site. **(C)** NIC203 (blue) and EG6180 (yellow) are two *C. tropicalis* wild strains that were originally isolated in the Caribbean Islands of Guadeloupe and Puerto Rico, respectively. **(D)** Whole-genome sequencing of the resultant strain from 19 generations of maternal backcross of NIC203 into the EG6180 background. Three separate loci maintain NIC203 alleles **(E)** developmental phenotypes in F_2_ progeny from the NIC203xEG6180 cross. 71% of progeny show a variety of developmental defects, including embryonic/larval lethality (Emb/L1 lethal), Larval arrest and Developmental delay (Dev. delay), which are in low frequency in the F_1_ or the parental strains **(F)** Genetic crosses with nearly isogenic lines (NILs) carrying a single NIC203 introgressed region confirm the presence of three independent maternal-effect TAs in Chr. II, III, and V. No significant developmental defects were observed in the parental NIL strains (top row). When a NIL is crossed to EG6180, ~25% of the F_2_ self-progeny are affected. Maternal backcrosses (third row) double the proportion of affected progeny, whereas parental backcross-progeny (fourth row) aren’t affected. Genotyping progeny (bottom bars) shows are found predominantly in homozygotes for the EG6180 genotype. Errors bars indicate 95% confidence intervals calculated with the Agresti-Coull method (*57*).

We previously discovered two TAs in the nematode *Caenorhabditis elegans*: the paternal-effect *peel-1/zeel-1* element (*8*, *9*) and the maternal-effect *sup-35/pha-1* element (*10*). The discovery of the latter was particularly unexpected because *pha-1*, the zygotic antidote, was long thought to be an organ-specific differentiation gene (*11*). These elements, together with the *Medea* element in the *Tribolium* flour beetle (*12*), are the only currently known TAs in animals. However, the possible mischaracterization of TAs as developmental genes, together with their more recent discovery in fungi and plants (*6*, *13*–*15*), led us to hypothesize that TAs may in fact be common, and that additional TAs could exist in other *Caenorhabditis* species. We tested this hypothesis in *Caenorhabditis tropicalis*, formerly known as *C. sp.* 11 (*16*, *17*). *C. tropicalis* is found exclusively in tropical regions worldwide and is distantly related to *C. elegans*; the genome sequences of the two species are as divergent as those of humans and mice (*18*). While most *Caenorhabditis* species have separate male and female sexes, both *C. elegans* and *C. tropicalis* have independently evolved hermaphrodites that self-fertilize but can also mate with males (Fig. 1B). Hermaphroditism has greatly facilitated the genetic study of *C. elegans* (*19*), and this advantage extends to *C. tropicalis*.

## Maternal-effect genetic incompatibility in *Caenorhabditis tropicalis*

A previous study reported that crosses between *C. tropicalis* isolates from different geographical locations are more likely to generate defective offspring than those between isolates from the same location (*17*). We were particularly intrigued by a cross between two strains isolated in the Caribbean, NIC203 (Guadeloupe, France) and EG6180 (Puerto Rico, USA) (Fig. 1C). Selfing of F_1_ progeny from the cross resulted in ~25% F_2_ embryonic lethality, which closely matches the expected proportion of affected progeny if a TA is present in one of the parents (*8*). We replicated this incompatibility between NIC203 and EG6180 (24.2% embryonic lethality, n=124; fig. S1) and found that it stems from an interaction between maternal and zygotic genotypes. In a cross between heterozygous F_1_ hermaphrodites and EG6180 males, the proportion of dead F_2_ embryos doubled (53.1%, n=207; fig. S1), whereas in the reciprocal cross between F_1_ males and EG6180 hermaphrodites, little lethality was observed (1.1%, n=182; fig. S1). This inheritance pattern indicates the presence of a maternal-effect TA in *C. tropicalis*, akin to the *sup-35/pha-1* TA in *C. elegans* and the *Medea* element in the beetle *Tribolium* (*10, 12*).

## Genome assembly of *C. tropicalis* strain NIC203

To enable the genetic study of the novel TA, we *de novo* assembled the genome of NIC203 using a combination of Illumina short reads and Oxford Nanopore long reads (Table S1). First, we generated three Nanopore runs (libraries fragmented to 8kb, 20kb, and unfragmented) and assembled the NIC203 genome into 21 scaffolds (N50 of 6,217,412bp, fig. S2A). To further improve the assembly, we sequenced the genomes of 350 F_2_ recombinant individuals from a NIC203 x EG6180 cross using a newly developed protocol (see Methods). We used the correlation between markers in the F_2_ sample to cluster and order the scaffolds into six linkage groups (fig. S2B), corresponding to the six *C. tropicalis* chromosomes. Synteny analysis supported our chromosome-level assembly, and allowed us to infer chromosome identity (fig. S2C). Lastly, we sequenced the transcriptome of NIC203 and created comprehensive gene annotations using both transcriptome-guided and *de novo* gene calling. Our final annotation includes 22,720 genes (24,833 transcripts), and analysis of conserved genes showed very high completeness (98.9% of conserved nematode genes were identified)

## Three distinct TAs in *C. tropicalis* strain NIC203

To map the locus underlying the novel maternal-effect TA, we took advantage of its inherent gene-drive activity (*20*). We backcrossed hybrid hermaphrodites to EG6180 males for 19 generations and sequenced the resulting strain (fig. S3). We expected the genetic background of the strain to be EG6180 with the exception of the region containing the putative NIC203 TA, which should have actively selected for its own presence throughout the backcross. To our surprise, we found not one but three regions carrying NIC203 alleles, which mapped to different chromosomes: II, III, and V (Fig. 1D). The same regions still carried NIC203 alleles after 30 generations of backcrossing, suggesting a strong selective force (fig. S4). We hypothesized that the two additional loci detected in our backcross were independent maternal-effect TAs that were missed in our initial phenotypic analysis because they did not induce embryonic lethality (fig. S1) but instead affected the fitness of larvae or adult worms. To test this hypothesis, we isolated F_2_ early embryos from a NIC203 x EG6180 cross and followed their individual growth at 24 h intervals for eight days. We found that a large fraction of the F_2_ progeny (71.2%, n=139) was affected by a wide range of defects, including not only embryonic lethality but arrest during larval growth and developmental delay (Fig. 1E). *C. tropicalis* embryos typically develop into adults and start laying eggs after two days at 25°C. Some affected F_2_ individuals, however, only developed to the second larval stage (L2) after two days, and started laying eggs after six or more days. Nevertheless, their progeny were viable and fertile. In sharp contrast, we did not detect significant levels of developmental defects in either parental strain (NIC203, 0%, n=115; EG6180, 3.4%, n=118) or in their F_1_ hybrid progeny (0%, n=31) (Fig. 1E). These observations are consistent with our hypothesis that the defects in the F_2_ progeny are the result of multiple TAs segregating in the cross.

To test whether multiple TAs are present in the cross, we generated nearly isogenic lines (NILs), each carrying only one of the NIC203 introgressed regions in the EG6180 background. We successfully obtained NILs for each of the three loci: NIL-Chr. II, NIL-Chr. III, and NIL-Chr. V. We confirmed the presence of only one introgressed region per strain and determined their boundaries by whole-genome sequencing (Table S2, fig. S5). We crossed each NIL to EG6180 and followed the phenotypes of F_2_ self-progeny, as well as progeny of maternal and paternal backcrosses. We found that for each NIL, the development of progeny homozygous for the EG6180 genotype was impaired (Fig. 1F). Furthermore, backcrossing F_1_ progeny from each cross to EG6180 revealed that the toxicity of all three TAs was transmitted maternally (Fig. 1F). The considerable phenotypic heterogeneity we observed in F_2_ progeny from the NIC203 x EG6180 cross reflects differences among the defects caused by these TAs: those on Chr. II and Chr. III cause developmental delay, while the one on Chr. V causes embryonic and L1 lethality. Although the NIC203 Chr. II TA is active at 25°C (fig. S6), we observed that the developmental delay is more pronounced at 20°C (Fig. 1F). None of these defects were observed in the parental NIL lines above background levels (Fig. 1F). In summary, *C. tropicalis* NIC203 carries three unlinked maternal-effect TAs, two of which, unlike all previously known TAs, do not induce embryonic lethality but instead cause larval arrest and developmental delay.

## Balancing selection of antagonizing TAs

Our NIL crosses indicate that the NIC203 maternal toxins are highly penetrant—most if not all of the progeny that don’t inherit them show abnormal phenotypes. Theoretically, three fully penetrant unlinked TAs should affect 57.8% of the F_2_ progeny (Fig. 2A). This prediction doesn’t fit the observed fraction of affected F_2_ progeny in the NIC203 x EG6180 cross (71.2%, n=139; Fig. 2A), suggesting that at least one additional TA could be segregating in this cross, perhaps contributed by the other parent, EG6180. To discover putative EG6180 TAs, we performed a multigenerational backcross of heterozygous F_1_ hermaphrodites to NIC203 males. In contrast to our previous backcross, this mating scheme is designed to specifically reveal EG6180 maternal-effect TAs. Whole-genome sequencing of the backcrossed strain after 32 generations revealed two EG6180 introgressed regions, on Chr. II and Chr. V (Fig. 2B). We generated two NILs, each carrying only one EG6180 introgressed region, mated the hermaphrodites from each NIL to NIC203 males, and phenotyped the F_2_ progeny. In addition, we performed maternal and paternal backcrosses for each NIL. The results show that both introgressed regions contain maternal-effect TAs which are absent from NIC203 (Fig. 2C). The EG6180 Chr. II TA causes developmental delay, whereas the one on Chr. V causes L2/L3 larval arrest/lethality (Fig. 2D). Neither phenotype was observed in the parental NIL lines (Fig. 2C).

**Figure 2.**
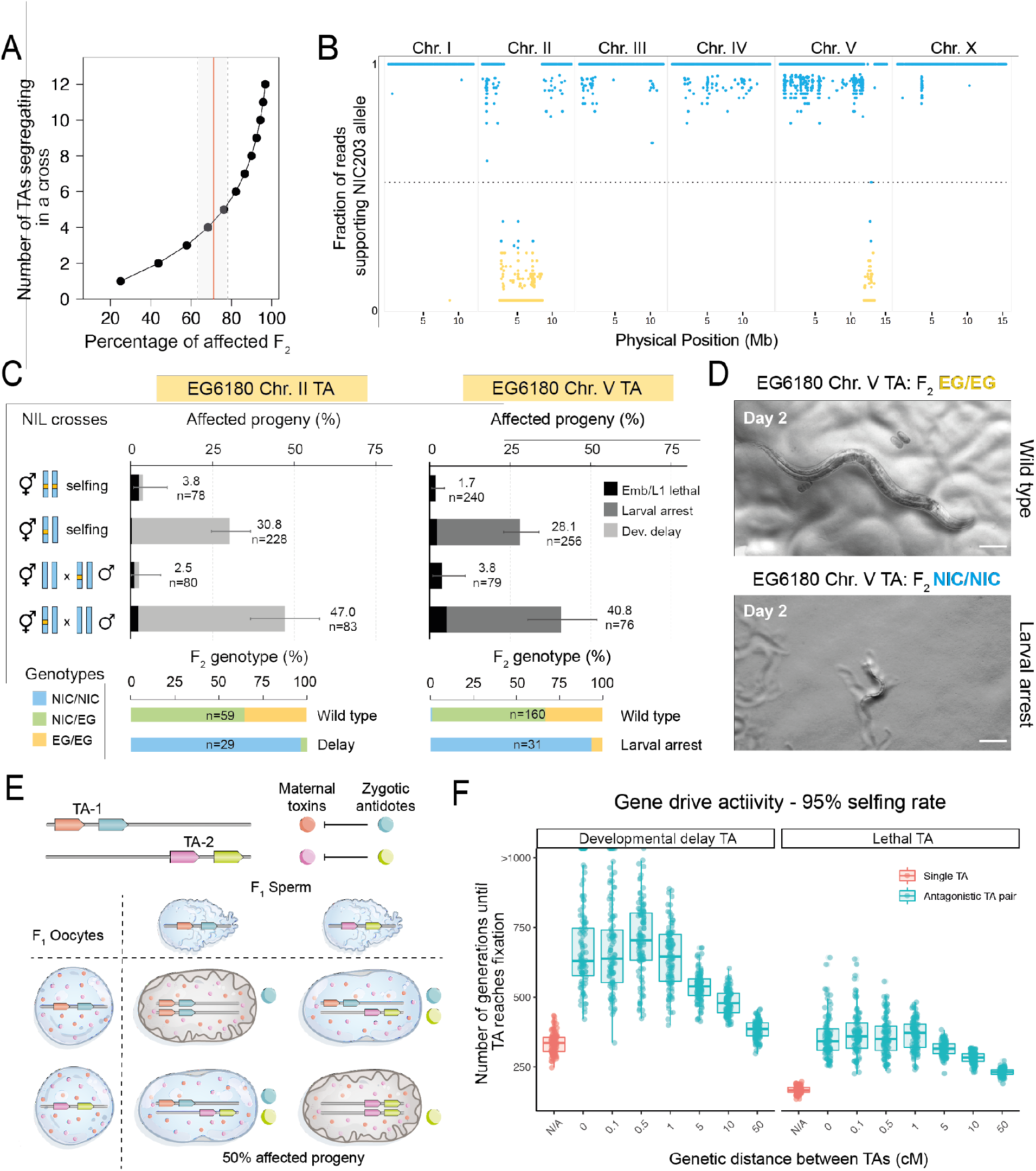
Balancing selection of antagonizing maternal-effect TAs. **(A)** The expected proportion of affected F_2_ progeny as a function of the number of unlinked TAs segregating in a cross between two isolates. A red vertical line depicts the observed proportion of affected progeny in the NIC203 x EG6180 cross (71.2%). The shaded area is a 95% CI. **(B)** Whole-genome sequencing of the resultant strain from 32 generations of maternal backcross of EG6180 into the NIC203 background reveals two regions, on Chr. II and Chr. V, carrying putative EG6180 TAs. **(C)** Crosses with nearly isogenic lines (NILs) carrying a single EG6180 introgressed region confirmed the presence of two independent maternal-effect TAs on Chr. II and V. No significant developmental defects were observed in the parental NILs. Genotyping shows affected progeny are predominantly homozygous for the NIC203 (NIC) genotype, while EG6180 homozygotes (EG) and heterozygotes (NIC/EG) are wild-type **(D)** The phenotype of worms affected by the Chr. V TA. Representative images of worms scored as phenotypically wild type (EG6180 homozygotes, top) or larval arrest (NIC203 homozygotes, bottom) after 2 days of development. Scale bar is 100 μm. **(E)** A model for antagonizing maternal-effect TAs. When two TAs are located on homologous chromosomes in close proximity, they counteract each other. In this case, two maternal toxins (salmon and magenta circles) are loaded into eggs prior to fertilization. Only heterozygous individuals express both zygotic antidotes (cyan and green circles). Punnett square depicts the expected percentage of affected F_2_ progeny for two fully penetrant TAs, up to 50% assuming no recombination between the TAs. For simplicity, only the toxins, and not the antidotes, are shown in the embryos. **(F)** We simulated the gene drive activity of single and antagonizing TAs. For each condition, we propagated 100 populations for 1,000 generations. Each population was seeded with 1,000 individuals, of which 500 were homozygous for each TA element. As a proxy to the androdioecious (hermaphrodite-male) mode of reproduction of *C. tropicalis*, we simulated a 95% selfing rate. We modeled TAs that induce developmental delay as TAs that decrease the fitness of non-carriers by half, compared to fully penetrant lethal toxins. We refer to a TA as fixed in the population once it reaches 0.95 allele frequency. Simulations with values >1,000 are those in which no TA reached fixation. For antagonizing TAs, we also modeled the genetic distance between pairs of TAs. 1 centiMorgan (cM) equals 1% meiotic recombination frequency.

Four of the five maternal-effect TAs are found as pairs on the same chromosome, with one present in NIC203 and the other in EG6180. The two Chr. V TAs are located on opposite arms, but those on Chr. II are relatively close to each other. We refer to such TA pairs, each contributed by one parent, as antagonizing TAs. Theoretically, in the absence of recombination between such closely located TAs, 50% of the F_2_ progeny should be affected, because only heterozygous individuals can counteract the effects of both toxins—they express both zygotic antidotes, whereas homozygotes express only one (Fig. 2E). Consequently, we would expect a large excess of heterozygotes among phenotypically wild-type worms. To test this prediction, we focused on the pair of TAs on Chr. II because they are located in the chromosome center, which has very low levels of recombination in nematodes (*21*). We performed a cross between NIC203 and EG6180 at 20°C, isolated F_2_ embryos, and followed their development for three days. Remarkably, genotyping of wild-type F_2_ individuals showed that 94.5% were NIC/EG heterozygotes for a marker on Chr. II:8.6Mb, in contrast to the expected 66.7% if the two toxins were segregating independently (n=55, χ²-test, p = 1.15×10-5).

In *C. elegans*, the *peel-1/zeel-1* and *sup-35/pha-1* TAs quickly spread in laboratory conditions due to their intrinsic gene drive activity (*8*, *10*, *22*). However, the observed pattern of heterozygous advantage in *C. tropicalis* Chr. II suggests that their gene drive activity is hindered. Over evolutionary time, this would result in a novel mechanism of balancing selection favoring heterozygous individuals and maintaining both TAs in wild populations. To better understand the effect of antagonizing TAs on gene drive dynamics, we simulated how single and antagonizing TA pairs spread in populations of hermaphroditic nematodes. We found that antagonizing TA pairs take significantly longer than single TAs to reach fixation, leading to more stable genetic incompatibilities (Fig. 3F). This effect was consistent across a wide range of parameters, including the genetic distance between TAs, their mode of action (causing embryonic lethality or partial reduction in fitness), and the selfing rate of the population (fig. S7). Overall, our results suggest that when in direct conflict, TAs can last longer in populations and lead to more stable incompatibilities due to balancing selection.

**Figure 3.**
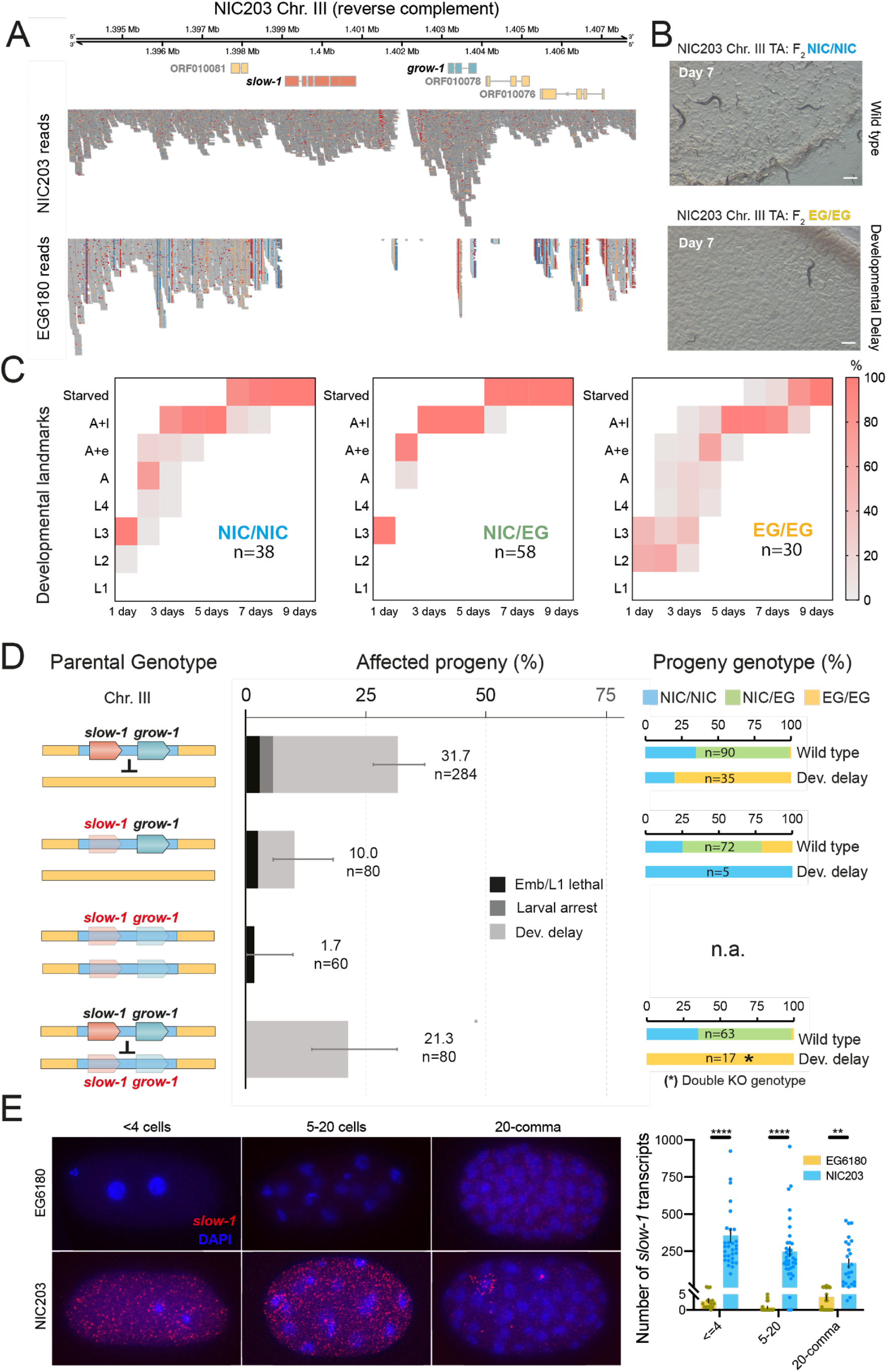
A TA causing developmental delay is encoded by *slow-1*, a toxin with homology to nuclear hormone receptors, and *grow-1*, it’s adjacent antidote. **(A)** An alignment of short reads from NIC203 (top) and EG6180 (bottom) to the NIC203 genome assembly in the Chr. III TA region shows four genes missing or divergent in EG6180. Mutations in two of the genes identified *slow-1* (salmon) and *grow-1* (blue) as the toxin and antidote genes. **(B)** Representative images of worms scored as phenotypically wild type (NIC203 homozygotes, top) and delayed (EG6180 homozygotes, bottom) 7 days after a single egg was seeded to the plate. Wild type plates already harbor F3 progeny, while the affected worm is arrested as a larva. Scale bar is 500 μm. **(C)** Developmental timing of the F_2_ progeny from a NIC203 Chr. III NIL x EG6180 cross for each genotype class. Single F_2_ worms seeded on 5cm plates were scored every 24 hours (L1-L4=larval stages, A=Adults, A+e=Adults that laid eggs, A+l=Adults and larvae, Starved=worms consumed all bacteria) **(D)** The genetic dissection of the toxin and antidote underlying the TA on NIC203 Chr. III. In the original NIL x EG6180 cross (top), EG6180 homozygous F_2_ progeny are delayed. A frameshift mutation in *slow-1* in the NIL background (2nd row) is sufficient to completely rescue the delayed phenotype of EG6180 homozygotes, indicating that *slow-1* is the underlying maternal toxin. An additional frameshift mutation in *grow-1* (3rd row), located immediately downstream of *slow-1*, does not cause any developmental defects by itself. However, the *slow-1 grow-1* double mutant is susceptible to the TA element, despite the strains being genetically identical apart from the two mutations. This shows that *grow-1* is the antidote to the *slow-1* toxin. All delayed progeny carry the double mutant genotype. **(E)** Representative images of NIC203 and EG6180 embryos hybridized with smFISH probes against *slow-1* (left). Embryos were grouped into three broad developmental categories based on their number of nuclei (DAPI staining) and the total number of slow-1 transcripts were quantified using an automatic pipeline (right). All tests are unpaired t-tests, p-values are Bonferroni corrected (** P ≤ 0.01, **** P ≤ 0.0001).

## A nuclear hormone-like maternal toxin causes developmental delay in *C. tropicalis*

Only two TAs have been genetically dissected in animals, and both kill embryos (*8*, *10*). We set out to identify the genes underlying the TA in Chr. III of NIC203 because the toxin induced a novel TA-associated phenotype—developmental delay (Fig. 1F). Alignment of Illumina short reads from EG6180 to our NIC203 assembly revealed four genes within the introgressed NIL region that were extremely divergent or missing in EG6180 (*ORF010076*, *ORF010078*, *ORF010079*, and *ORF010080*; Fig. 3A). To identify the toxin, we knocked out each of these genes in the NIC203 Chr. III NIL with CRISPR/Cas9 (*23*). We then crossed WT and mutant NIL hermaphrodites to EG6180 males and characterized their F_2_ progeny. In the WT cross, F_2_ progeny homozygous for the EG6180 genotype at the introgressed region typically took five days to start laying eggs, whereas TA carriers require only two (Fig. 3B and 3C). Mutations in two of the genes (*ORF010076* and *ORF010078*) did not impair the ability of the TA to cause developmental delay (fig. S8). However, a frameshift mutation in the second exon of *ORF010080* was sufficient to drastically reduce the percentage of delayed and arrested worms from 28.9% to 7.5%, indicating that *ORF010080* encodes the toxin (n=139, Fig. 3D and fig. S9). *ORF010080* encodes a 366-amino-acid protein that has significant homology to nuclear hormone receptors (NHR) (fig. S10). We named this gene *slow-1* (for maternal-effect SLOWer development). As expected for a knock-out of the maternal toxin, all homozygous EG/EG F_2_ individuals in this cross were phenotypically wild type (n=15, Fig. 3D). The residual fraction of developmentally delayed F_2_ worms (7.5%) were all NIC/NIC homozygotes, suggesting that there is an additional and independent factor, likely contributed by EG6180, that affects NIC203 individuals with a low penetrance.

Finally, we determined that *ORF010079*, which we named *grow-1* (for zygotic rescue of GROWth), is the zygotic antidote. To show this, we first knocked out *grow-1* with CRISPR/Cas9 in the background of the *slow-1* mutant (fig. S8). Next, we crossed *slow-1 grow-1* double-mutant hermaphrodites to the parental Chr. III NIL and inspected the F_2_ progeny. If these two genes encode the toxin and the antidote, we would expect the double mutant to behave like EG6180, which lacks both genes (Fig. 3A). Indeed, we observed that 21.3% (n=80) of the F_2_ progeny were delayed, and all developmentally delayed individuals were homozygous for the *slow-1 grow-1* double-mutant allele (Fig. 3D). Moreover, we observed background levels of abnormal phenotypes in the *slow-1* or the *slow-1 grow-1* knockout parental strains, suggesting that these two genes do not play any significant role in the normal development of the worm (Fig. 2D and fig. S11). In summary, because loss of *slow-1* abrogates the toxicity of the TA, and mutations in both *slow-1* and *grow-1* are sufficient to turn the TA allele into an EG6180-like susceptible one, we conclude that *slow-1* and *grow-1* are the underlying toxin and antidote, respectively.

Like most members of the NHR family, *slow-1* contains a predicted ligand-binding domain; however, two properties set it apart from most of its homologs: it lacks a DNA binding domain, and it contains two transmembrane domains on its C-terminus (fig. S10). *grow-1*, on the other hand, encodes a novel small protein—only 125 amino acids long—that has no homology or predicted protein domains. In agreement with its role as a maternal-effect toxin, we detected *slow-1* mRNA expression in NIC203 embryos prior to zygotic gene activation (<4 cell stage) using single-molecule *in situ* hybridization (Fig. 2E). As expected, we did not detect *slow-1* expression in EG6180 embryos (Fig. 2E). *slow-1* transcripts decreased during embryonic development and were undetectable at the comma stage, with the exception of a single pair of cells, which likely correspond to germline precursors Z2 and Z3 based on their position in the embryo. This observation suggests that the toxin is transcribed in the germline as early as during embryogenesis.

## A maternal-effect TA in *C. briggsae*

Our discovery that some TAs in *C. tropicalis* induce developmental delay prompted us to re-evaluate a previously described genetic incompatibility in *C. briggsae* (*24*), the other known hermaphroditic relative of *C. elegans* (Fig. 1B). Crosses between the reference strain AF16 (Ahmedabad, India) and HK104 (Okayama, Japan) result in a significant fraction of developmentally delayed F_2_ individuals carrying markers for the AF16 genotype on Chr. III (*24*). However, the mechanism underlying this incompatibility is unknown. We hypothesized that the delay could be caused by a maternal-effect TA present in HK104. To test this hypothesis, we crossed HK104 to AF16, selfed F_1_ hermaphrodites, and backcrossed F_1_ males and hermaphrodites to AF16 (Fig. 4). These crosses replicated the incompatibility between AF16 and HK104: 20.1% (n=214) of the F_2_ self-progeny showed developmental delay, while no delay was observed in either parental strain. Furthermore, 40 of the 41 delayed F_2_ individuals were homozygous for the AF16 genotype at the previously reported incompatibility region on Chr. III. Backcrossing F_1_ hermaphrodites to AF16 males doubled the proportion of affected progeny (39,0%, n=159), while no delay was observed in the reciprocal backcross (0%, n=70, respectively, Fig. 4). These results indicate that the *C. briggsae* isolate HK104 carries a maternal-effect TA that induces developmental delay, analogous to the *slow-1/grow-1* TA in *C. tropicalis*. Overall, our results demonstrate that TAs are common among *Caenorhabditis* species.

**Figure 4.**
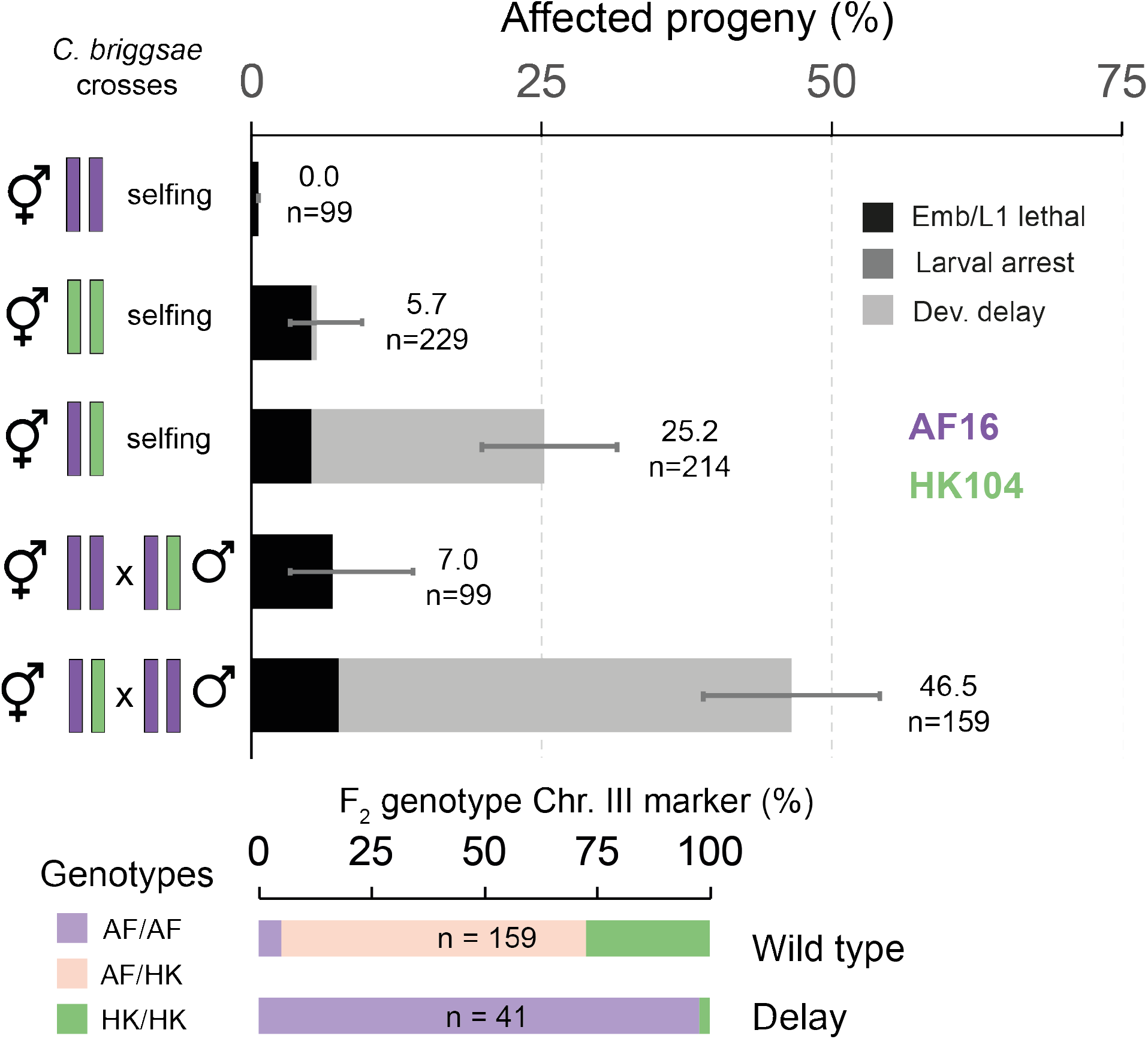
A maternal-effect TA in *C. briggsae* causes developmental delay. Crosses between two *C. briggsae* isolates, AF16 (purple) and HK104 (green), recapitulate a previously reported genetic incompatibility (*24*). Developmentally delayed individuals are observed among their F_2_ progeny but not in the parental strains. As expected from previous work, 97.6% of delayed individuals are AF16 homozygotes in a genetic marker in the Chr. III incompatibility region, whereas wild type progeny are heterozygous or HK104 homozygotes. We found that the delay phenotype follows the expected pattern of inheritance for a maternal-effect TA, with the proportion of delayed individuals doubled in the maternal backcross while no delayed individuals are observed in the paternal backcross. Embryonic lethality observed in the crosses is consistent with the lethality in the HK104 parental strain. Errors bars indicate 95% confidence intervals calculated with the Agresti-Coull method.

## Discussion

We discovered five maternal-effect TAs in *C. tropicalis* and one TA in *C. briggsae*. Together with our previous work in *C. elegans*, we have identified TAs in all known hermaphroditic *Caenorhabditis* species. We showed that the mode of action of TAs is not restricted to targeting meiosis, as in fungi and plants (*6*, *13*–*15*), or disrupting embryonic development (*8*, *10*, *12*), but that, surprisingly, they can also affect larval and adult stages. TAs that exclusively target post-embryonic development have been largely overlooked in nematodes and other model organisms. We identified the genes underlying a novel TA, *slow-1/grow-1*, which delays the larval development of progeny that do not inherit it. Future work will focus on understanding the molecular mechanism by which *slow-1* induces developmental delay. We hypothesize that *slow-1* acts as a dominant-negative peptide, binding a small lipophilic molecule, which normally acts as a NHR ligand, and sequestering it to a membrane. In support of this view, mutations in some conserved NHR genes such as *daf-12* and *nhr-25* are known to cause heterochronic phenotypes in *C. elegans* and to affect larval and larval-to-adult transitions (*25*, *26*). The NHR family is massively expanded and rapidly evolving in *Caenorhabditis* nematodes (*27*). SLOW-1 only has very weak homology to other NHR family members in *C. tropicalis*— it is most similar to NHR-47 (17.9% identity and 30.2% similarity)—suggesting that it has strongly diverged from its likely NHR ancestor (fig. S9).

Balancing selection preserves allelic diversity in populations in the face of genetic drift (*28*, *29*). We discovered a new mechanism by which balancing selection can arise—conflict between antagonizing TAs. In this scenario, heterozygous individuals are fitter than homozygotes not because they are better adapted to their environment, but simply as a by-product of conflict between selfish elements. Each TA selectively targets individuals homozygous for the other element and only heterozygotes are protected (Fig. 2E). This phenomenon could occur in any diploid organism and is reminiscent of lab-derived balancer chromosomes that maintain deleterious mutations in the heterozygous state (*30*). Importantly, antagonizing TAs provide an effective way to decouple genetic incompatibilities from gene drive. Genetic incompatibilities stemming from a single TA are transient; they disappear once the selfish element reaches fixation in the population. However, antagonizing TAs result in more stable incompatibilities (fig. S7), which could create barriers to genetic exchange between populations. Balancing selection of antagonizing TAs could explain the paradoxical retention of TAs in selfing species and, more generally, perhaps the existence of regions of extreme nucleotide divergence among wild isolates (*31*, *32*). All TAs known to date are found within hyper-divergent regions (*8*, *10*, *32*). We hypothesize that TAs originated in the gonochoristic ancestors of extant hermaphroditic species and have been maintained in their genomes via their antagonism. TAs have independently evolved in bacteria, archaea, plants, and fungi (*33*). Our discovery of these elements in *C. elegans* (*8*, *10*) and now in *C. tropicalis* and *C. briggsae* strongly suggests that TAs are common in nematodes and raises the question of whether they are also present in other animal groups, including vertebrates.

## Acknowledgements

We thank members of the Burga and Kruglyak labs for their comments. E.B-D. is supported by NIH Grant K99-HG010369, CB is supported by the Centre National de la Recherche Scientifique (CNRS), A.B. is supported by the Austrian Academy of Sciences and the European Research Council (ERC-2019-StG-851470), and L.K. is Supported by the Howard Hughes Medical Institute and NIH grant R01 HG004321. E.B-D., A.B., and L.K. wrote the manuscript; all authors discussed and agreed on the final version of the manuscript. The authors declare no competing financial interests. Sequencing data are available under NCBI bioproject PRJNA649308.

## Material and Methods

### *C. tropicalis* and *C. briggsae* maintenance, strains, and mutant alleles

All experiments were carried out at 25°C unless specified in the text. *C. tropicalis* and *C. briggsae* strains were grown on modified nematode growth medium (NGM) containing *E. coli* OP50, and 1% agar/0.7% agarose to prevent burrowing of wild isolates using standard culturing techniques developed for *C. elegans* (*19*). A list of all the strains used and generated in this study is available at Table S3. Some of the strains were provided by the CGC, which is funded by the NIH Office of Research Infrastructure Programs (P40 OD010440).

### Multigenerational maternal backcrosses for mapping TAs

To map putative TAs, we took advantage of their intrinsic gene drive activity. We reasoned that by performing maternal backcrosses (using heterozygous mothers) for multiple generations, the genomic regions containing TAs could be identified by whole-genome sequencing because they would be refractory to the backcross. Furthermore, the more generations of backcross, the smaller the expected sizes of the introgressed regions. To map NIC203 TAs, we mated NIC203 hermaphrodites to EG6180 males, picked ~5 L4 F_1_ hermaphrodites and mated them with a 5-fold excess EG6180 males. Then, we took ~5 L4 F_2_ hermaphrodites and mated them again with EG6180 males. In total we repeated this cycle 19 times (resulting strain is QX2340; fig. S3). Although we expect most of the L4 progeny originate from a mating event, we cannot exclude the possibility that some may be the product of selfing. To identify EG6180 maternal-effect TAs, we performed an analogous multigenerational backcross for 34 generations but mated heterozygous hermaphrodites to NIC203 males (resulting strain is INK74).

### Generation of nearly-isogenic lines (NILs)

To generate NILs carrying only a single maternal-effect TA, we took as a starting point the multigenerational maternal backcross strains QX2340 and INK74. In the case of NIC203 TAs, we first crossed QX2340 hermaphrodites (carrying NIC203 regions in Chr. II, III and V in an otherwise EG6180 background) to EG6180 males. Next, we took F_1_ males and mated them with EG6180 hermaphrodites. Using heterozygous F_1_ males (in contrast to hermaphrodites) guarantees the retrieval of viable/healthy EG/EG non-carriers among their progeny because the toxins are not transmitted via sperm. We isolated and genotyped F_2_ hermaphrodites at the three introgressed loci and selected those that were heterozygous NIC/EG for one TA and homozygous EG/EG non-carriers for the other loci (set of primers used available in table S4). Finally, we genotyped F_3_ hermaphrodites to generate homozygous NIL strains for each TA: QX2341 (NIC Chr. II), QX2345 (NIC Chr. III), and QX2342 (NIC Chr. V). In the case of EG6180 TAs, crosses were performed in an analogous fashion but using as a starting point INK74 hermaphrodites (carrying EG6180 regions in Chr. II, and V in an otherwise NCI203 background) and NIC203 males. The resulting NILs strains were: INK75 (EG Chr. II) and INK77 (EG Chr. V). Set of primers used available in table S4. The genotype of all five NILs was confirmed by whole-genome Illumina short-read sequencing (fig. S5).

### Illumina short-read whole-genome sequencing

For sequencing NIC203, EG6180, INK74, INK75 and INK77, we extracted genomic DNA (gDNA) using the DNeasy Blood & Tissue kit (Qiagen) following manufacturer protocol with the exception that worms were frozen and thawed to facilitate lysis prior to DNA extraction. gDNA for the rest of the strains was extracted using the Masterpure Complete DNA and RNA purification kit (Lucigen) using the protocol for tissue samples, with the modification that worms were frozen and thawed to facilitate lysis prior to extraction. We prepared Illumina sequencing libraries using Nextera (Illumina), Nextera flex (Illumina), or a homebrew modified Tn5 enzyme (see the Single-worm sequencing section). Table S1 includes information about the sequencing libraries generated for each strain. Libraries were quantified using the Qubit HS kit and sequenced on an Illumina Nextseq, Illumina Hiseq X or Hiseq 4000 instruments.

### Nanopore long-read sequencing

We extracted high molecular weight (HMW) genomic DNA using the Masterpure Complete DNA and RNA purification kit (Lucigen), using the protocol for tissue samples, with the modification that worms were frozen and thawed to facilitate lysis prior to extraction. We then prepared 8 kb, 20 kb and unfragmented sequencing libraries using the 1D Ligation Sequencing Kit from Oxford Nanopore Technologies (SQK-LSK109). 8 kb and 20 kb fragmentation was done using g-TUBE (Covaris) following manufacturer protocols. Libraries were loaded on a MinION MK1B device (Oxford Nanopore). Read calling was done using MinKNOW software.

### *De novo* genome assembly

We carried out a hybrid assembly strategy inspired by a previous study with modifications (*34*). First, we normalized Illumina short reads from NIC203 to 100X average coverage using BBNorm (sourceforge.net/projects/bbmap) with default parameters. We then assembled short reads using the MaSuRCA genome assembler (*35*), adjusting Jellyfish hash size based on an estimated genome size of 80 Mb. Long reads were then error corrected and trimmed using Canu (*36*). We used the short read assembly, together with the short reads, to correct long reads using Halc (*37*). Halc-corrected reads were then input to assembly using the Flye assembler with default parameters, setting estimated genome size to 80 Mb (*38*). Initial assembly included 32 scaffolds, with N50 of 4,750,223bp The assembly was error corrected using Hypo using long and short reads as input. We further scaffolded the polished assembly to 23 scaffolds using LRScaf (*39*). 2 short scaffolds were removed, since they aligned to *E. coli*, likely reflecting residual bacterial contamination in the Nanopore preps, resulting in 21 scaffolds with N50 of 6,217,412bp.

### Single-worm sequencing

We developed a novel protocol for whole-genome sequencing of single worms. Our protocol uses a modified version of the Tn5 transposase, an enzyme that forms the basis of the Illumina Nextera library prep kits. A homebrew version of the Nextera library prep protocol was previously published (Picelli et al) (*40*). We acquired plasmid pTXB1-Tn5 from Addgene and used the QuikChange Lightning Site-Directed Mutagenesis Kit (Agilent) and the Q5 site-directed mutagenesis kit (NEB) to recreate 4 amino acid changes that were previously found to improve the robustness of tagmentation to lower DNA concentrations (*41*). We then purified the modified Tn5 protein as described in Picelli et al, with the modification that we concentrated raw protein eluates using Amicon Ultra 15 Centrifugal Filter (MilliporeSigma) and size selected on a Superdex 200 Gel Filtration Column (GE-Healthcare). Since Tn5 only dimerizes in contact with DNA (*42*), we selected for the Tn5 monomer (50 kDa). Dialysis was done as described in Picelli et al. Tn5 aliquots were stored directly after purification in −80**°**C. A working aliquot was thawed and supplanted with glycerol and loaded with transposons, as described in Picelli et al. Working stock concentration was 1 μM.

For single-worm library prep, we sought to minimize volumes while carrying all steps in one tube. Each worm was suspended in 21 active units (0.035 μl) Proteinase K (Qiagen), 1 μl 5x TAPS (see Picelli et al for formulation) and 2.5 μl water, to a total of 3.5 μl (to ensure accurate amounts of proteinase K, a large mix was premade). Worms were lysed for 30 minutes in 65**°**C, followed by 5 minutes inactivation of proteinase K in 95**°**C. The inactivated lysate was then topped with 0.5 μl dimethyl formamide (DMF), 0.1 μl charged TN5 and 0.9 μl water. The tagmentation reaction was incubated for 7 minutes in 55**°**C. Tn5 was stripped by adding 1.25 μl 0.2% SDS, and incubated for an additional 7 minutes in 55**°**C. Library prep PCR was then done by adding 6.75 μl KAPA HiFi MasterMix (Roche), 0.75 μl DMSO and 0.75 μl EvaGreen (Biotium). PCR was done in an Agilent AriaMX qPCR machine, for 17-18 cycles, until reaction appeared to plateau. To reduce cross contamination, unique dual indexes (UDI) were used (barcodes were based on Illumina Cat no. 20027213 and were custom ordered from IDT). 2 μl from the final qPCR reaction were visualized on a gel for quality control, then each plate of 96 individually indexed worms were pooled and gel purified together selecting for 500 bp. Purified libraries were sequenced on Illumina Hiseq X.

### Single nucleotide variant calling

Variant calling was done using the *variant2* pipeline implemented in the *bcbio-nextgen* ngs data analysis software suite (https://github.com/bcbio/bcbio-nextgen). Reads from NIC203 and EG6180 were aligned to the 21 scaffold genome assembly using *BWA-mem* (*43*) and variant calling was done using *Genomic Analysis Toolkit (GATK Ver 4)* (*44*) as well as *Strelka* (*45*). Only variants that were identified by both tools were kept. Both tools were run with a ploidy setting of 2. Variants were filtered to include only those called as homozygous reference in NIC203 and homozygous alternative in EG6180. These variants were then used for inferring linkage groups using single-worm F_2_ sequencing (see below). The process was repeated after generating the final assembly to retrieve a finalized list of variants that include 105,431 single-nucleotide variants (SNVs) and 13,234 indels.

### Assembly to chromosome level using F2 single-worm genotyping

We aligned sequences from 350 F2 worms to the 21 scaffold genome using BWA-mem. We then called genotypes in each of the curated variants between NIC203 and EG6180 using *GATK haplotypecaller*. The genotyping in those individuals was then imputed using LB-impute (*46*), taking a constant 1 Mb interval as equivalent to 50% recombination chance in the absence of a genetic map, which isn’t available in *C. tropicalis*. Our single-worm sequencing resulted in very low coverage, and only the 254 individuals had imputed genotype information in at least half of the markers. To aid in linkage group computation, the imputed genotypes were then passed through a greedy filtering algorithm that removed each consecutive variant if the correlation between it and the previous one was over 0.95 across all individuals with a non-missing genotype. This resulted in 2,547 variants. r/qtl was then used to identify 6 linkage groups (Fig S2B). Due to the high completeness of the original assembly, each linkage group only included 2-4 scaffolds. The filtering process was repeated with a more stringent cutoff of 0.99 keeping a total of 7,356 variants. The variants within each linkage group were then ordered using R package ASMap (*47*) to determine the orientation and order of the scaffolds.

For synteny analysis, the NIC203 genome draft was aligned against *C. elegans* WBcel235 reference genome using Cactus (*48*, *49*). The aligned file was converted to the *maf* format, deduplicated using *Maftools* (*50*) and converted to links format using *awk*. Alignments were read into *R*, and consecutive alignments that were separated by less than 10,000bp were joined. Finally, alignments were plotted using the *Circos* R package (*51*) to generate the synteny plot in Figure S2.

### Genome annotation

We used Repeatmodeler to generate consensus repeat families for NIC203, followed by Repeatmasker to soft-mask repeats in the genome. A total of 9,057,221bp (11.10%) of the genome was masked. We generated RNA-seq libraries from mixed-stage populations of NIC203 worms using the TruSeq Stranded mRNA kit (Illumina) following manufacturer protocol, and sequenced them on a 2×75 paired-end Hiseq 3000 lane. The NIC203 soft-masked genome assembly and RNA-seq reads were input to *funannotate*, a pipeline for genome annotation (https://funannotate.readthedocs.io/). *Funannotate* is mostly used for annotating fungal genomes, so we used the following recommended modifications for annotating a nematode genome: we allowed an intron size of up to 20,000bp (based on the largest introns in *C. elegans*), changed *Busco* database to *nematoda*, and specified the *-species “other”* flag where recommended in the online documentation of the pipeline.

### Phenotyping and genotyping of parental and NIL TAs crosses

We set up crosses in 12-well plates. Typically, 5 hermaphrodites were mated with at least 5-fold excess males. After 3 days, ~10 F_1_ L4 hermaphrodites were placed into individual 5 cm plates. On day 4, once F_1_ hermaphrodites were fully gravid, they were transferred into fresh 5-cm plates and allowed to lay eggs for 4-5 hours. Then, we collected ~10 F_2_ embryos per F_1_ hermaphrodites. Each embryo was placed in a single 5 cm plate, which allowed us to reliably score post-embryonic phenotypes. Once F_2_ embryos were collected, we genotyped their mothers, and only included in our final analysis true F_1_ mothers and their progeny and not those that originated from self-fertilization of the original hermaphrodite. We scored the phenotype of F_2_ progeny every 24 hours typically for 5-9 days depending on the cross. Plates that initially contained a single phenotypically WT F_2_ hermaphrodite starved after 6-7 days. Following phenotyping, we genotyped each worm by PCR using indels (Table S4). Maternal and maternal backcrosses were scored in an analogous way. For maternal backcrosses, we did not include in our final calculations data from hermaphrodites that did not mate (no males were observed among their progeny) or hermaphrodites with a low outcrossing rate (<20% of progeny were males). Male progeny from maternal backcrosses for NIC203 Chr. II and NIC203 Chr. III TAs were included in final calculations of affected progeny. That is, we could clearly distinguish developmental delay in males. In the case of EG6180 Chr. II TA, we only included hermaphrodites due to uncertainty in male phenotyping. To study the genotype of NIC203 x EG6180 F_1_ progeny, we used a EG6180 strain carrying an spontaneous dumpy recessive mutation. We crossed WT NIC203 males to EG6180 dpy hermaphrodites and phenotyped only F_1_ heterozygous WT progeny.

### Fine-mapping of NIC203 Chr. III TA and CRISPR/Cas9 genome editing

The NIL strain carrying the NIC203 Chr. III TA (QX2345) carries a small ~40 kb introgression (Chr. II: 10,426,642-10,466,762 Mb). Within this introgression, we noticed a region of high divergence between NIC203 and EG6180 that mainly encompassed four predicted genes based on our RNA-seq guided annotation: *ORF010076*, *ORF010078*, *ORF010079*, and *ORF010080* (Fig. 2A). These four genes are located next to each other in the genome. We reasoned these genes were good candidates for a TA pair because toxin and antidotes are usually found in tight genetic linkage and are missing or highly mutated in the susceptible strain. Our model predicts that a knockout of the putative toxin should be perfectly tolerated by the parental carrier strain but should lead to a loss of the F_2_ incompatibility when crossed to the susceptible strain. We generated putative loss of function alleles for three candidate genes in this region (*ORF010076*, *ORF010078*, and *ORF010080*) in the genetic background of the NIL Chr. III. To generate *C. tropicalis* mutants we adapted the CRISPR/Cas9 ribonucleoprotein (RNP) complex injection protocol developed by Dokshin et al for *C. elegans* (*52*). In brief, we incubated 15 pmol of Cas9 (0.5 μl of Alt-R Cas9 from IDT) with 4.45 nM modified synthetic single guide RNA (sgRNA) (Synthego) for 10 minutes in 37℃ for 10 minutes. To the assembled RNP we then added 2.5 ng/μl of pCFJ90 plasmid as a co-injection marker and injected young adult hermaphrodites. For *ORF010079* we also included 2.2 ng/μl of a single-stranded oligonucleotide donor (ssODN) repair template that introduced a stop-codon and a frameshift. sgRNAs and repair template sequences can be found in Table S4. We singled F_1_ mCherry positive fluorescent worms and screened their progeny for random frameshift deletions or homology-directed repair events by Sanger sequencing. We then crossed each of these NIL mutants to EG6180 males. Of these three genes, only mutations in *ORF010080* abolished the delay observed among the homozygous EG/EG F_2_ progeny. We renamed *ORF010080 slow-1*. Since loss of function mutations in the putative antidote would result in sick or abnormal worms, we generated a knockout in the putative antidote *ORF010079* in the background of the mutant toxin. In the absence of the toxin, the antidote should be dispensable. Furthermore, alleles with mutations in both the toxin and the antidote should behave like susceptible EG6180 alleles in crosses. We first generated a knockout allele of *ORF010079* (the gene immediately downstream of the toxin) in the *grow-1* mutant genetic background. The targeted mutation incorporates a premature stop codon in the first exon. We then crossed the wild type NIC Chr. III NIL to the double mutant NIC Chr. III NIL. As expected from knocking out the antidote, all the F_2_ progeny that were homozygous for the double mutant allele were developmentally delayed. We named the antidote *ORF010079* as *grow-1*.

### Single molecule in situ hybridization

We designed 48 Custom Stellaris FISH Probes against *slow-1* mRNA using the Stellaris RNA FISH Probe Designer (Biosearch Technologies, Inc., Petaluma, CA) available online at www.biosearchtech.com/stellarisdesigner. NIC203 and EG6180 embryos were hybridised with *slow-1* probes labeled with Quasar® 570 (Biosearch Technologies, Inc.) following the manufacturer’s instructions. We did not perform smFISH on *grow-1*, due to the very small size of the transcript; the total number of fluorescent probes that could hybridize to the mRNA was only 15 (25 is the minimum recommended). The smFISH protocol was adapted from Raj et al (*53*). Embryos were imaged on an Axio Imager.Z2 (Carl Zeiss) widefield microscope equipped with a Hamamatsu Orca Flash 4 camera. Images were acquired using a 63x/1.4 plan-apochromat Oil DIC objective with the following filters: for DAPI Ex 406/15nm, Em 457/50nm and for Quasar® 570 Ex 545/30nm, Em 610/75nm. We acquired for each image a Z-stack containing 40 slices with a step size of 0.2 um. Quantification was performed using a Matlab script from the Raj lab available at https://bitbucket.org/arjunrajlaboratory/rajlabimagetools/

### *C. briggsae* maternal-effect incompatibility

We studied a previously described genetic incompatibility in a cross between the reference *C. briggsae* AF16 strain from India and the wild isolate HK104 from Japan (Table S3) (*24*). In their original study, Ross et al. reported that approximately 20% of the F_2_ progeny exhibited a pronounced developmental delay at 20°C. This phenotype was strongly associated with a genetic marker on Chr. III that also showed transmission ratio distortion in RIL hybrid lines. Delayed individuals were AF/AF homozygous. Importantly, delayed worms were not observed in the parental lines nor F_1_ progeny. This inheritance pattern suggested was compatible with the presence of TA in HK104. Interestingly, Baird et al. reported that the Chr. III associated incompatibility had a maternal-effect. When the authors backcrossed heterozygous mothers to AF16 males, they observed 10-25% delayed worms among their progeny but no delay was observed in the reciprocal paternal backcross (*54*). However, for a maternal-effect TA, we would expect approximately ~40-50% delayed progeny in the maternal backcross (two times the proportion of delayed individuals in F_1_ selfing) and not 10-25%. We hypothesized that the low levels of incompatibility reported in the maternal backcross experiments by Baird et al. could originate from the usage of sperm-depleted hermaphrodites for genetic crosses. In fact, it was later shown that sperm depleted *C. briggsae* hermaphrodites are still able to self-fertilize due to incomplete purging of self-sperm (*55*). To test this hypothesis, we crossed AF16 and HK104 parental lines at 25°C, performed maternal and paternal backcrosses and carefully genotyped each F_1_ and F_2_ progeny discarding any progeny that resulted from self-fertilization. Using our more stringent criteria, we observed 39% delayed progeny in the maternal backcross and no delay in the paternal backcross, as expected for a maternal-effect TA element. Primers used for genotyping are found in Table S4. Males from maternal backcrosses were not included in final counts.

### Gene drive simulations

Simulations were performed using the SLiM evolutionary framework (v 3.4) (*56*). A population was seeded with 500 individual carriers of TA1 and 500 carriers of TA2. A non Wright-Fisher simulation was carried out for 1,000 generations. A population size limit of 10,000 was enforced by scaling the fitness of each individual by 10,000. In each generation, a custom *modifyChild* function was used to simulate the action of a TA. Since sexes were not explicitly defined in the model, one parent was taken to be the mother. Offspring who did not inherit the TA element from a carrier mother were removed from the population. Penetrance was implemented as a probabilistic modifier for killing (killing of affected offspring would occur at probability *p* = penetrance). We used penetrance of the TA as a proxy to study TAs that do not kill progeny prior to egg laying but only cause developmental delay, decreasing the fitness of non-carriers. We also simulated a wide range of genetic distances between antagonizing TAs to understand the impact of recombination. For each set of parameters, we simulated 100 populations. To run the simulation across many different parameters, we wrote an R script to run SLiM from the command line. We plotted the number of generations required until at least one TA reached fixation in the population, to be able to directly compare the rate of one TA to antagonistic TAs. We defined a TAs as fixed once it reached 0.95 allele frequency. The script for the simulation is available at github: https://github.com/eyalbenda/TA_simulations/

## Supplementary Material

### Supplementary tables

**Table S1.**
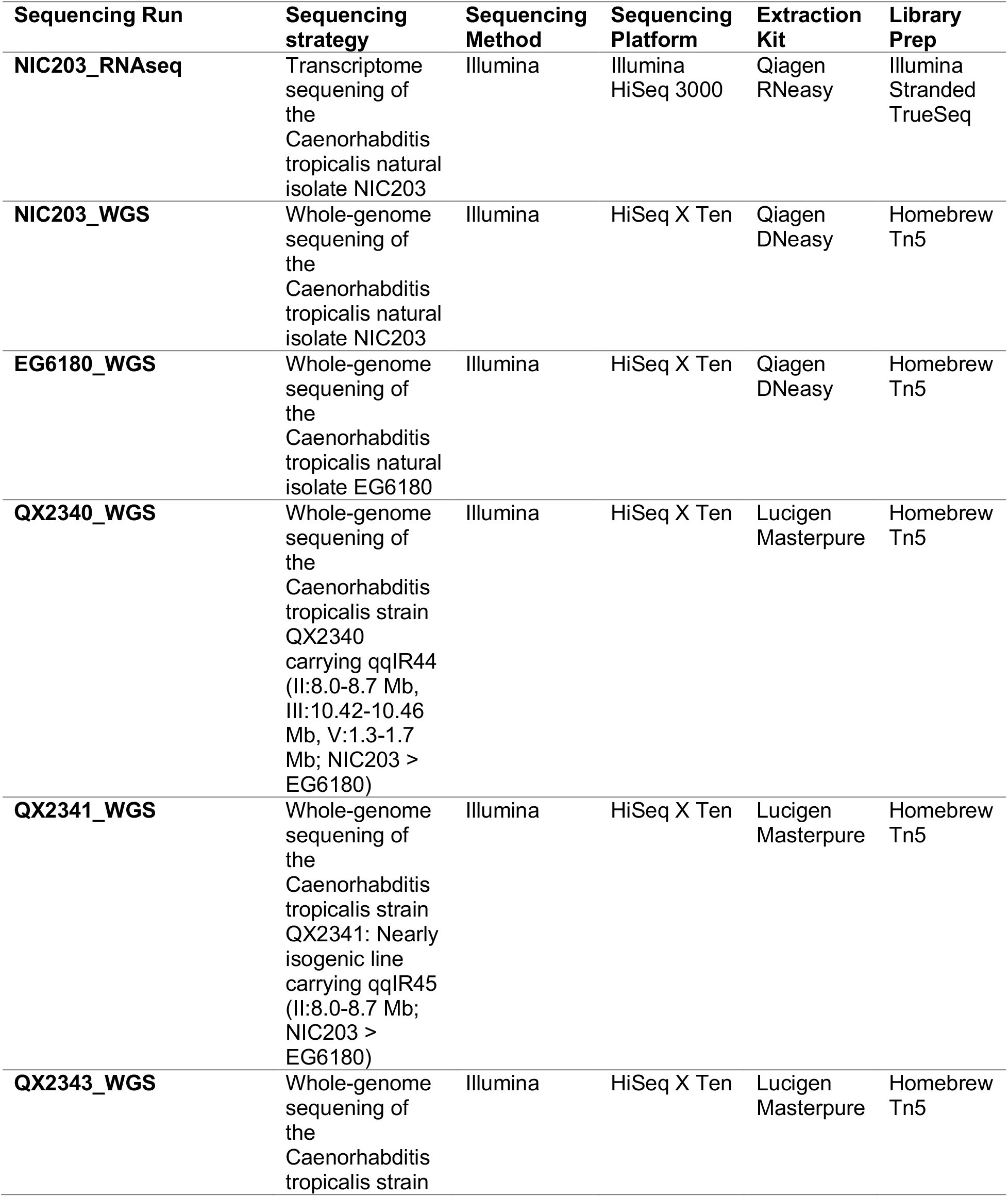
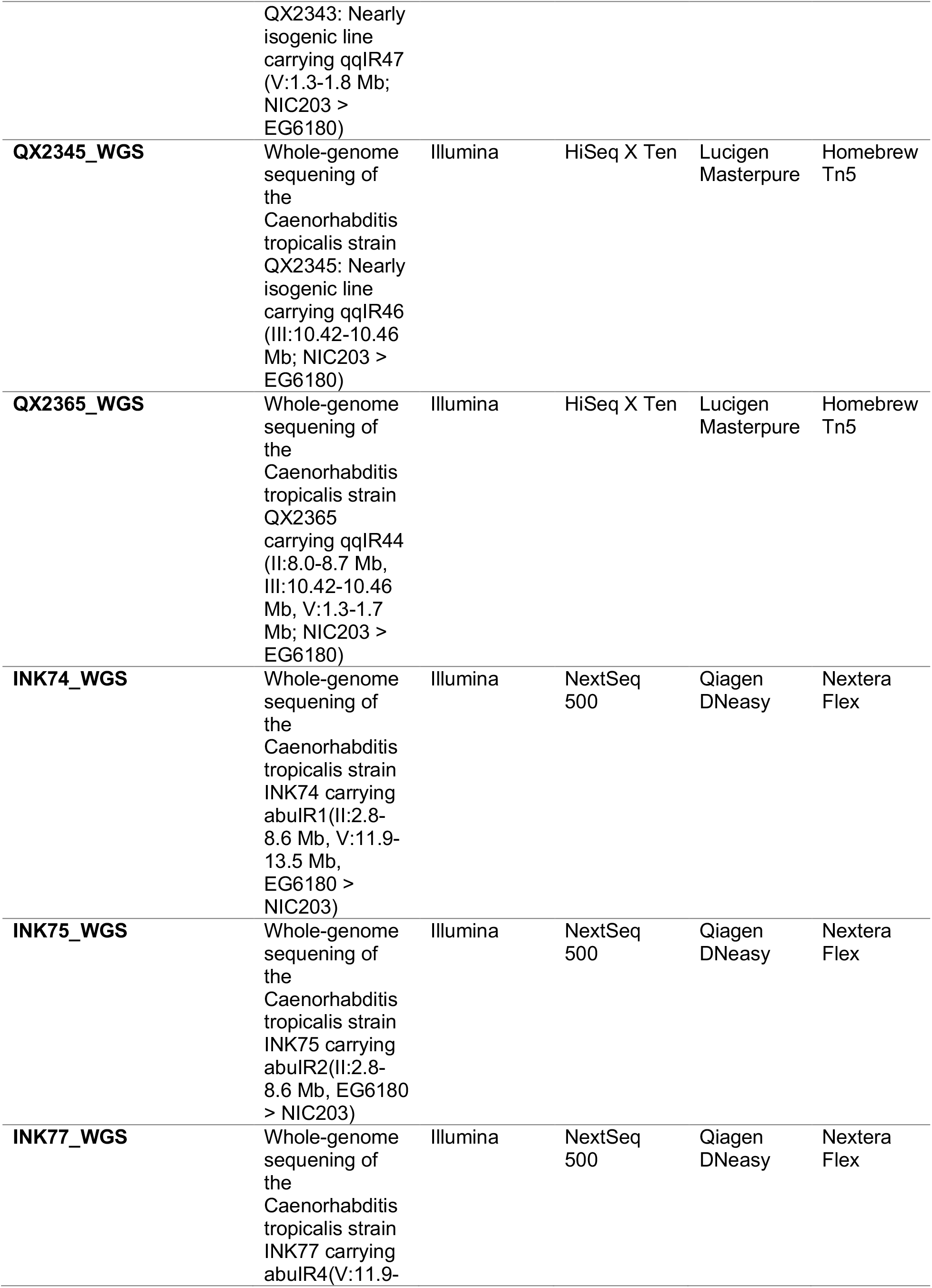
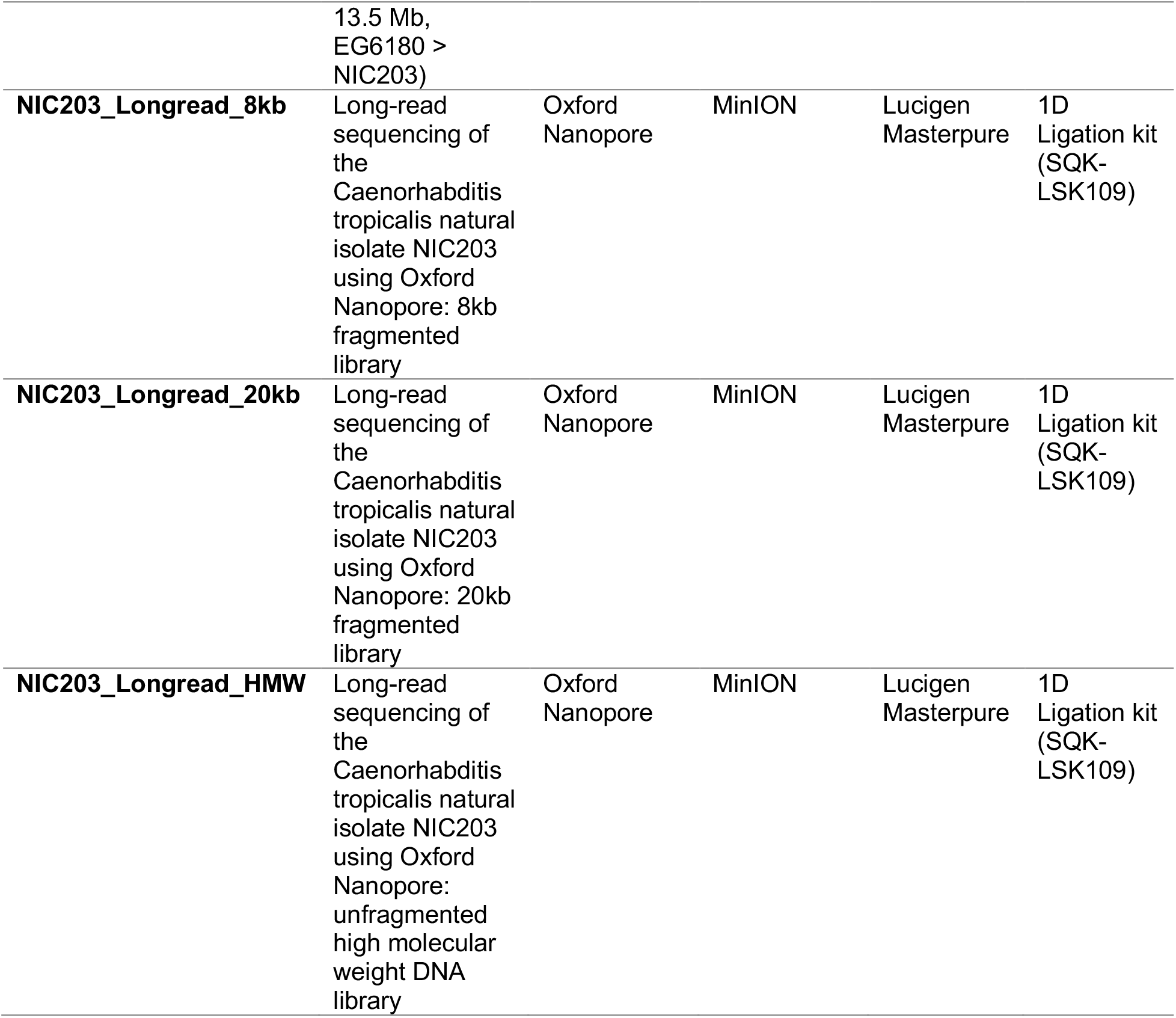
Summary of Illumina and Nanopore sequencing libraries.

**Table S2.** Genomic coordinates of introgressions in NILs carrying TAs.

| **NIL** | **Introgression**<br>**Start (bp)** | **Introgression**<br>**End (bp)** | **Introgression**<br>**Length (bp)** | **TA**<br>**element** | **Phenotype** |
| --- | --- | --- | --- | --- | --- |
| **NIC203**<br>**Chr. II TA** | 8,009,743 | 8,756,516 | 746,773 | Maternal-effect | Dev. delay |
| **NIC203**<br>**Chr. III TA** | 10,426,642 | 10,466,762 | 40,120 | Maternal-effect | Dev. delay |
| **NIC203**<br>**Chr. V TA** | 1,289,235 | 1,751,145 | 461,910 | Maternal-effect | Emb/L1<br>lethal |
| **EG6180**<br>**Chr. II TA** | 2,836,704 | 8,581,670 | 5,744,966 | Maternal-effect | Dev. delay |
| **EG6180**<br>**Chr. V TA** | 11,935,356 | 13,453,526 | 1,518,170 | Maternal-effect | Larval<br>arrest |

**Table S3.**
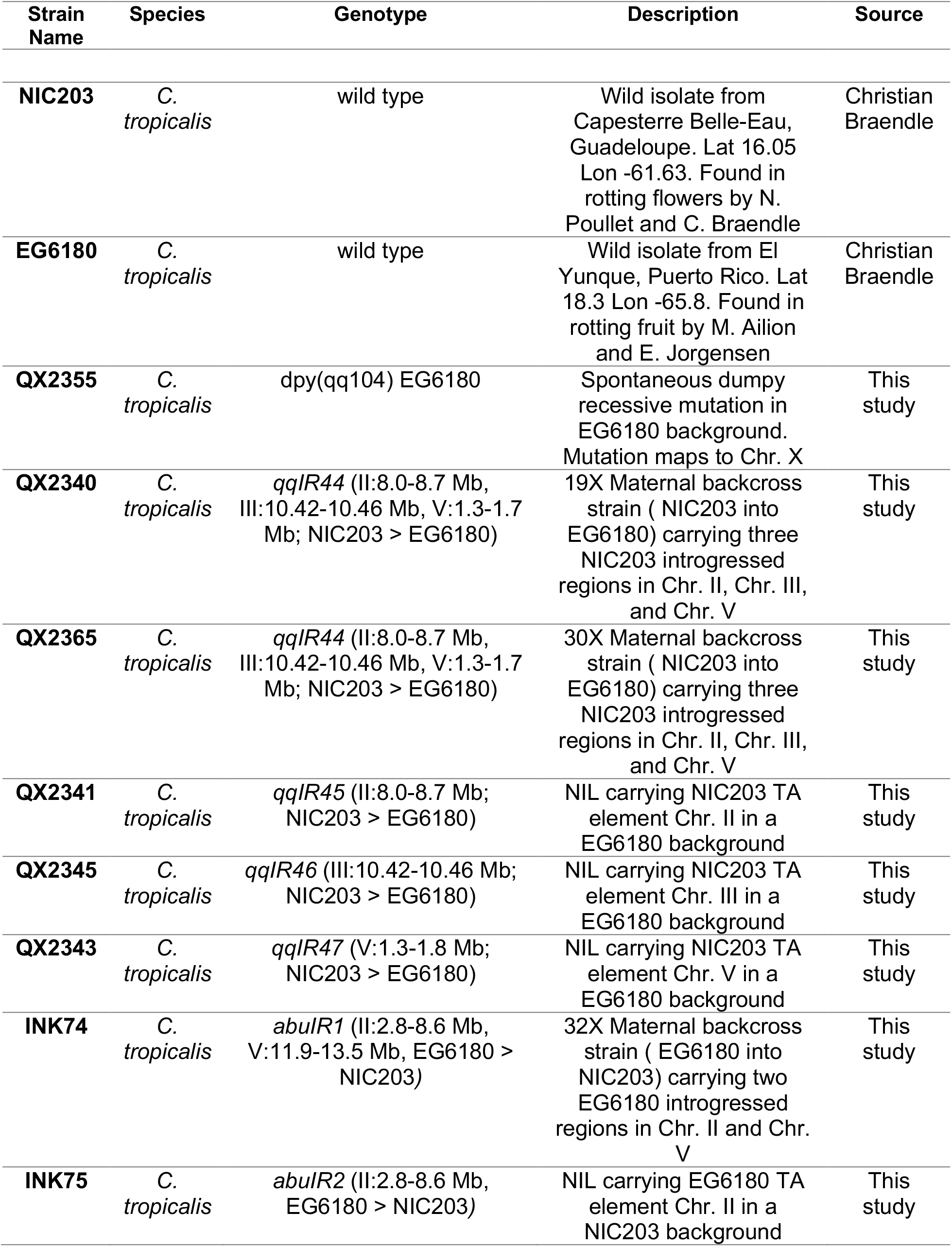
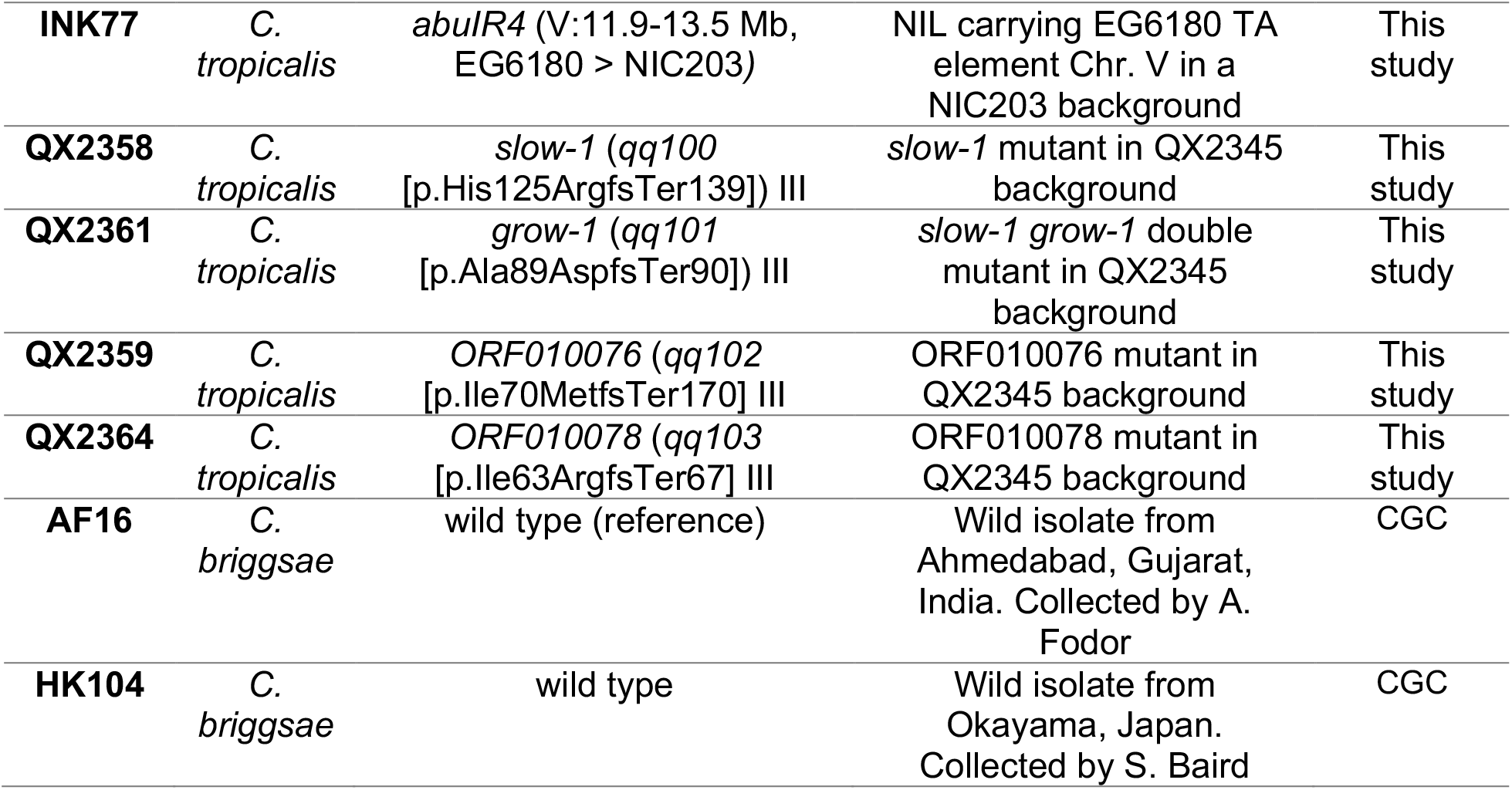
Strains used in this study.

| Strain Name | Species | Genotype | Description | Source |
| --- | --- | --- | --- | --- |
| **NIC203** | *C. tropicalis* | wild type | Wild isolate from Capesterre Belle-Eau, Guadeloupe. Lat 16.05 Lon -61.63. Found in rotting flowers by N. Pouillet and C. Braendle | Christian Braendle |
| **EG6180** | *C. tropicalis* | wild type | Wild isolate from El Yunque, Puerto Rico. Lat 18.3 Lon -65.8. Found in rotting fruit by M. Ailion and E. Jorgensen | Christian Braendle |
| **QX2355** | *C. tropicalis* | dpy(qq104) EG6180 | Spontaneous dumpy recessive mutation in EG6180 background. Mutation maps to Chr. X | This study |
| **QX2340** | *C. tropicalis* | *qqIR44* (II:8.0-8.7 Mb, III:10.42-10.46 Mb, V:1.3-1.7 Mb; NIC203 > EG6180) | 19X Maternal backcross strain ( NIC203 into EG6180) carrying three NIC203 introgressed regions in Chr. II, Chr. III, and Chr. V | This study |
| **QX2365** | *C. tropicalis* | *qqIR44* (II:8.0-8.7 Mb, III:10.42-10.46 Mb, V:1.3-1.7 Mb; NIC203 > EG6180) | 30X Maternal backcross strain ( NIC203 into EG6180) carrying three NIC203 introgressed regions in Chr. II, Chr. III, and Chr. V | This study |
| **QX2341** | *C. tropicalis* | *qqIR45* (II:8.0-8.7 Mb; NIC203 > EG6180) | NIL carrying NIC203 TA element Chr. II in a EG6180 background | This study |
| **QX2345** | *C. tropicalis* | *qqIR46* (III:10.42-10.46 Mb; NIC203 > EG6180) | NIL carrying NIC203 TA element Chr. III in a EG6180 background | This study |
| **QX2343** | *C. tropicalis* | *qqIR47* (V:1.3-1.8 Mb; NIC203 > EG6180) | NIL carrying NIC203 TA element Chr. V in a EG6180 background | This study |
| **INK74** | *C. tropicalis* | *abulR1* (II:2.8-8.6 Mb, V:11.9-13.5 Mb, EG6180 > NIC203) | 32X Maternal backcross strain ( EG6180 into NIC203) carrying two EG6180 introgressed regions in Chr. II and Chr. V | This study |
| **INK75** | *C. tropicalis* | *abulR2* (II:2.8-8.6 Mb, EG6180 > NIC203) | NIL carrying EG6180 TA element Chr. II in a NIC203 background | This study |
| **INK77** | *C.*<br>*tropicalis* | *abulR4* (V:11.9-13.5 Mb,<br>EG6180 > NIC203) | NIL carrying EG6180 TA<br>element Chr. V in a<br>NIC203 background | This<br>study |
| **QX2358** | *C.*<br>*tropicalis* | *slow-1* (*qq100*<br>[p.His125ArgfsTer139]) III | *slow-1* mutant in QX2345<br>background | This<br>study |
| **QX2361** | *C.*<br>*tropicalis* | *grow-1* (*qq101*<br>[p.Ala89AspfsTer90]) III | *slow-1 grow-1* double<br>mutant in QX2345<br>background | This<br>study |
| **QX2359** | *C.*<br>*tropicalis* | *ORF010076* (*qq102*<br>[p.Ile70MetfsTer170]) III | ORF010076 mutant in<br>QX2345 background | This<br>study |
| **QX2364** | *C.*<br>*tropicalis* | *ORF010078* (*qq103*<br>[p.Ile63ArgfsTer67]) III | ORF010078 mutant in<br>QX2345 background | This<br>study |
| **AF16** | *C.*<br>*briggsae* | wild type (reference) | Wild isolate from<br>Ahmedabad, Gujarat,<br>India. Collected by A.<br>Fodor | CGC |
| **HK104** | *C.*<br>*briggsae* | wild type | Wild isolate from<br>Okayama, Japan.<br>Collected by S. Baird | CGC |

**Table S4.**
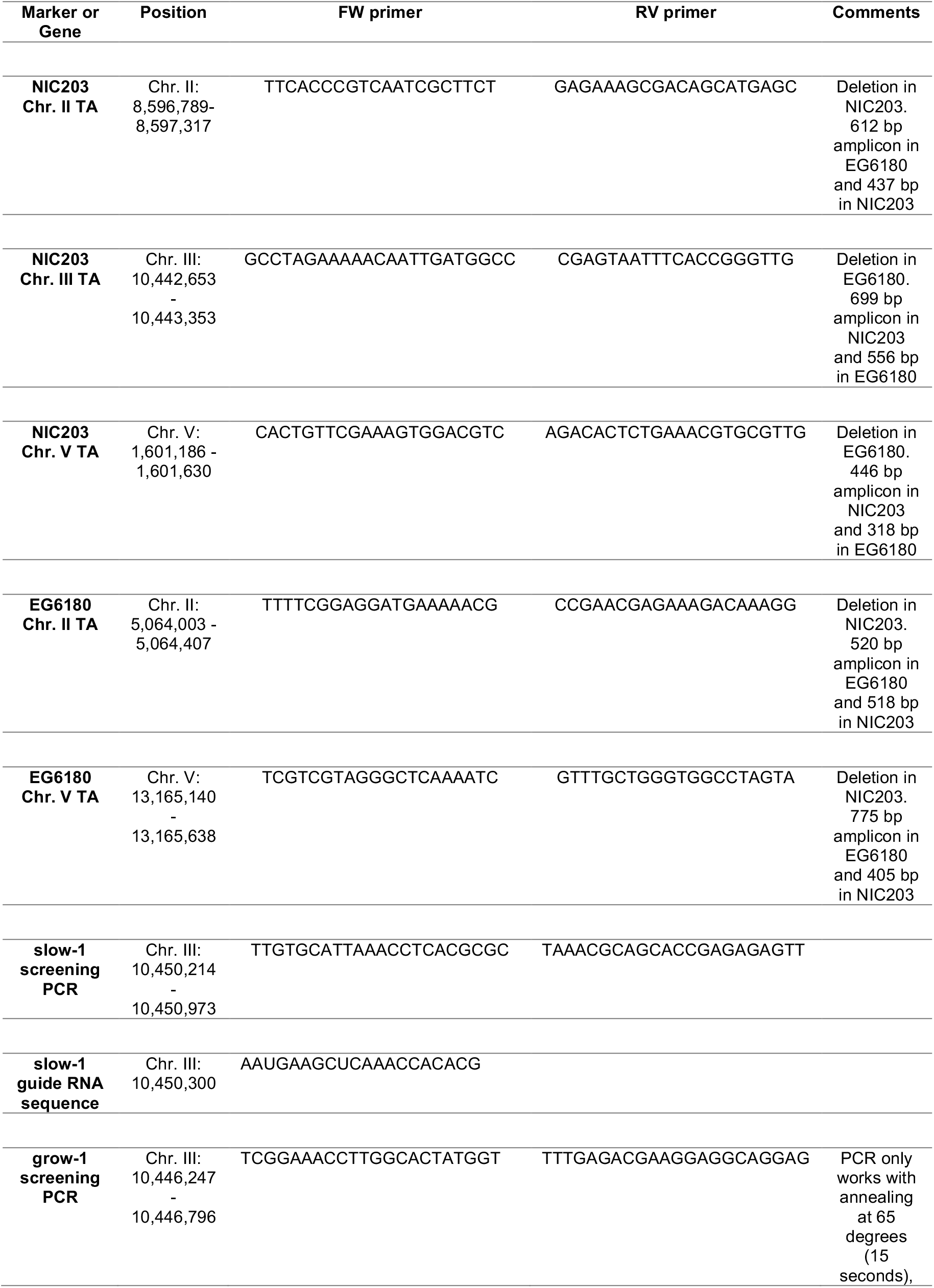
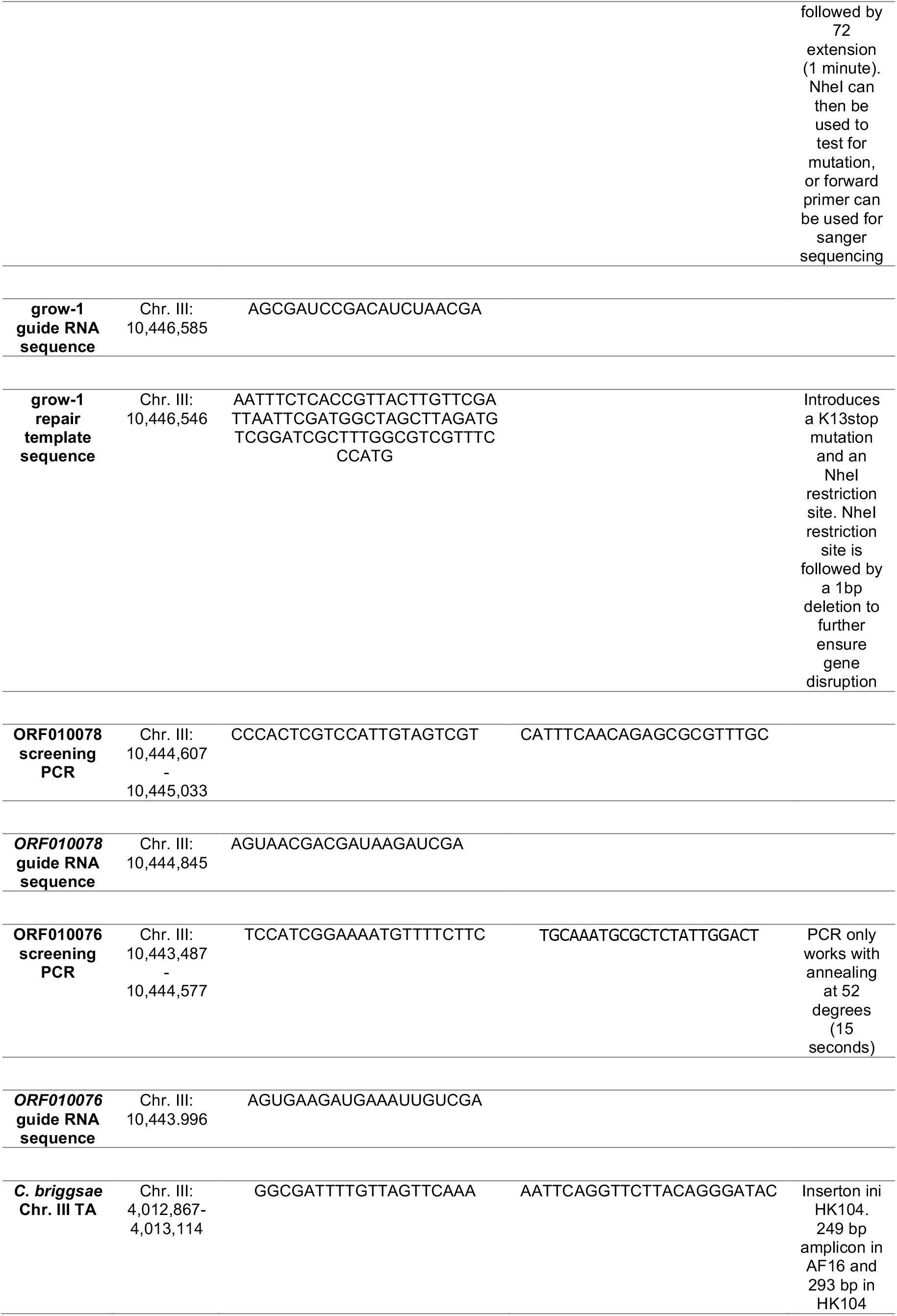
Primers and oligos used in the study.

### Supplementary Figures

**Figure S1.**
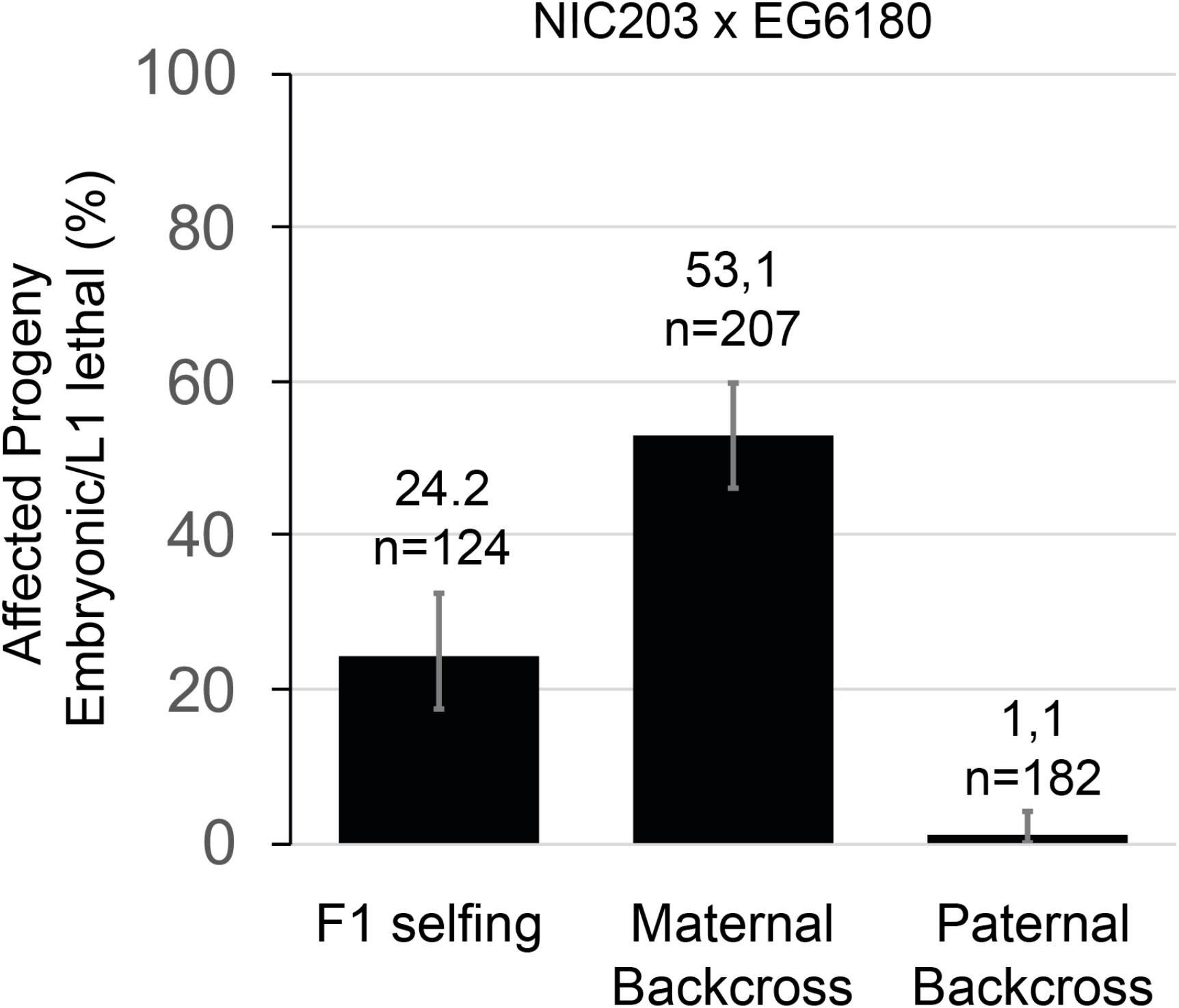
A maternal factor causes embryonic and L1 lethality among F_2_ progeny of a cross between NIC203 and EG6180 wild isolates. We observed ~25% embryonic and L1 lethality among the F2 progeny of selfing heterozygous hermaphrodites. The proportion of dead progeny doubled among the progeny of a maternal backcross — heterozygous NIC/EG F1 hermaphrodites crossed to EG/EG males. The reciprocal cross, EG/EG hermaphrodites crossed with heterozygous NIC/EG F1 males only show background levels of lethality. These results are consistent with the presence of a maternal-acting TA element present in NIC203 and missing in EG6180. Errors bars indicate 95% confidence intervals calculated with the Agresti-Coull method.

**Figure S2.**
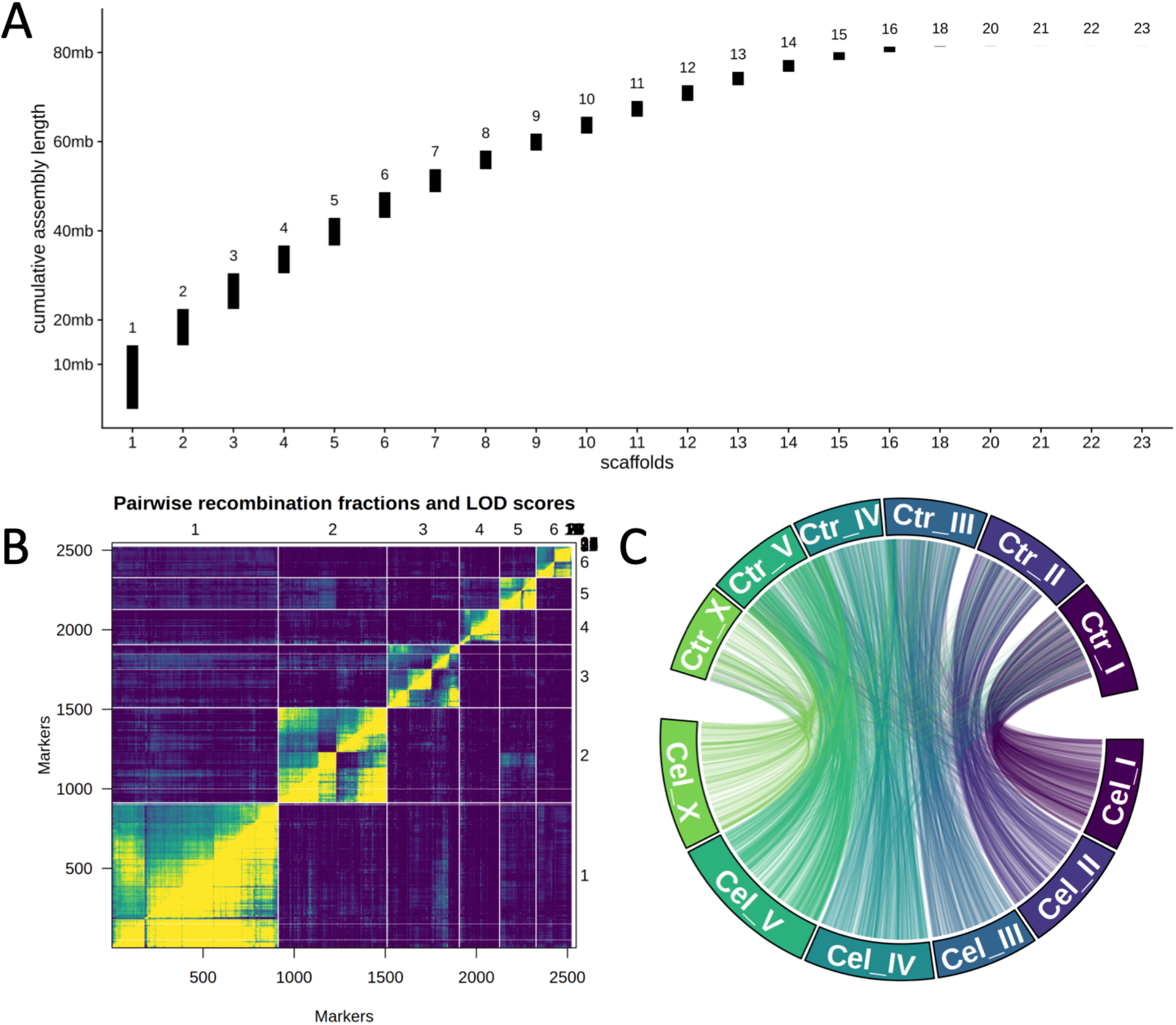
Chromosome-level assembly of the *C. tropicalis* NIC203 wild isolate strain. **(A)** Cumulative lengths of the 23 scaffolds in the original assembly. **(B)** Clustering of markers to linkage groups based on recombination patterns in the F_2_ progeny. **(C)** Synteny analysis between the NIC203 genome and the *C. elegans* genome identifies chromosomes.

**Figure S3.**
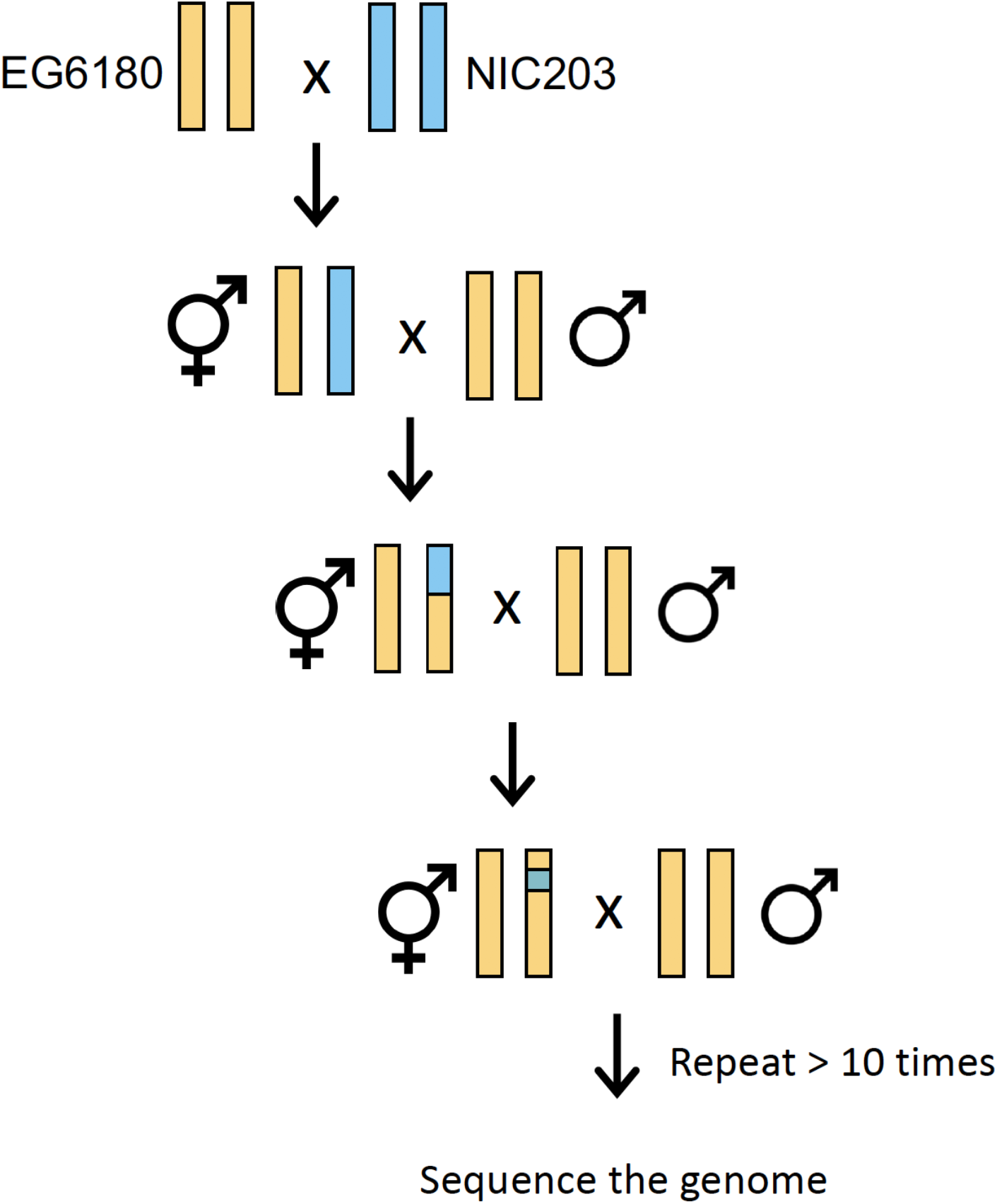
Scheme of the multigenerational backcross strategy to map maternal-effect TAs. For simplicity, only mapping of NIC203 materna-effect TAs is shown and only one chromosome is depicted. We initially crossed heterozygous NIC/EG mothers (~5 hermaphrodites) to >5-fold excess EG/EG males. If the NIC allele carries a maternal-effect TA element and the EG does not, then we would expect the TA to be active. This cross will result in NIC/EG (non-affected) and EG/EG (affected) individuals. For multiple generations, we randomly selected ~5 L4 hermaphrodites among the fastest developing progeny (likely enriched in NIC/EG individuals) and crossed them with EG/EG males. NIC203 genomic regions carrying putative TAs are expected to be retained despite extensive backcrossing and get smaller every generation due to recombination. Finally, we sequenced the whole genome of the resulting population and identified the introgressed regions.

**Figure S4.**
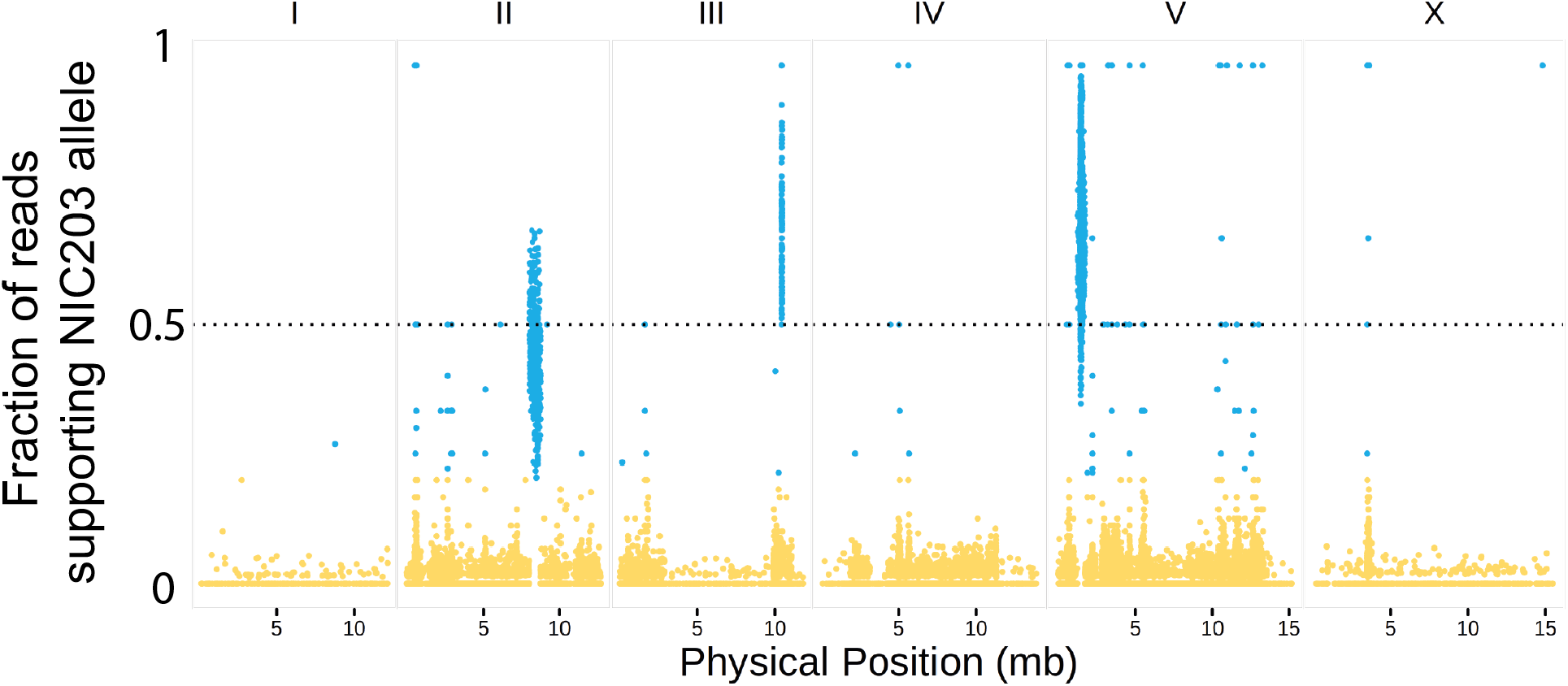
NIC203 regions carrying TAs are retained after 30 generations of maternal backcrossing. We performed a maternal backcross to map NIC203 TAs as shown in Figure S3. We sequenced the whole genome of the resulting population after 30 generations of backcross using Illumina short-read sequencing. Three genomic regions located in Chr. II, Chr. III, and Chr. V retained NIC203 alleles, indicating a strong selection in favor of the NIC203 haplotype.

**Figure S5.**
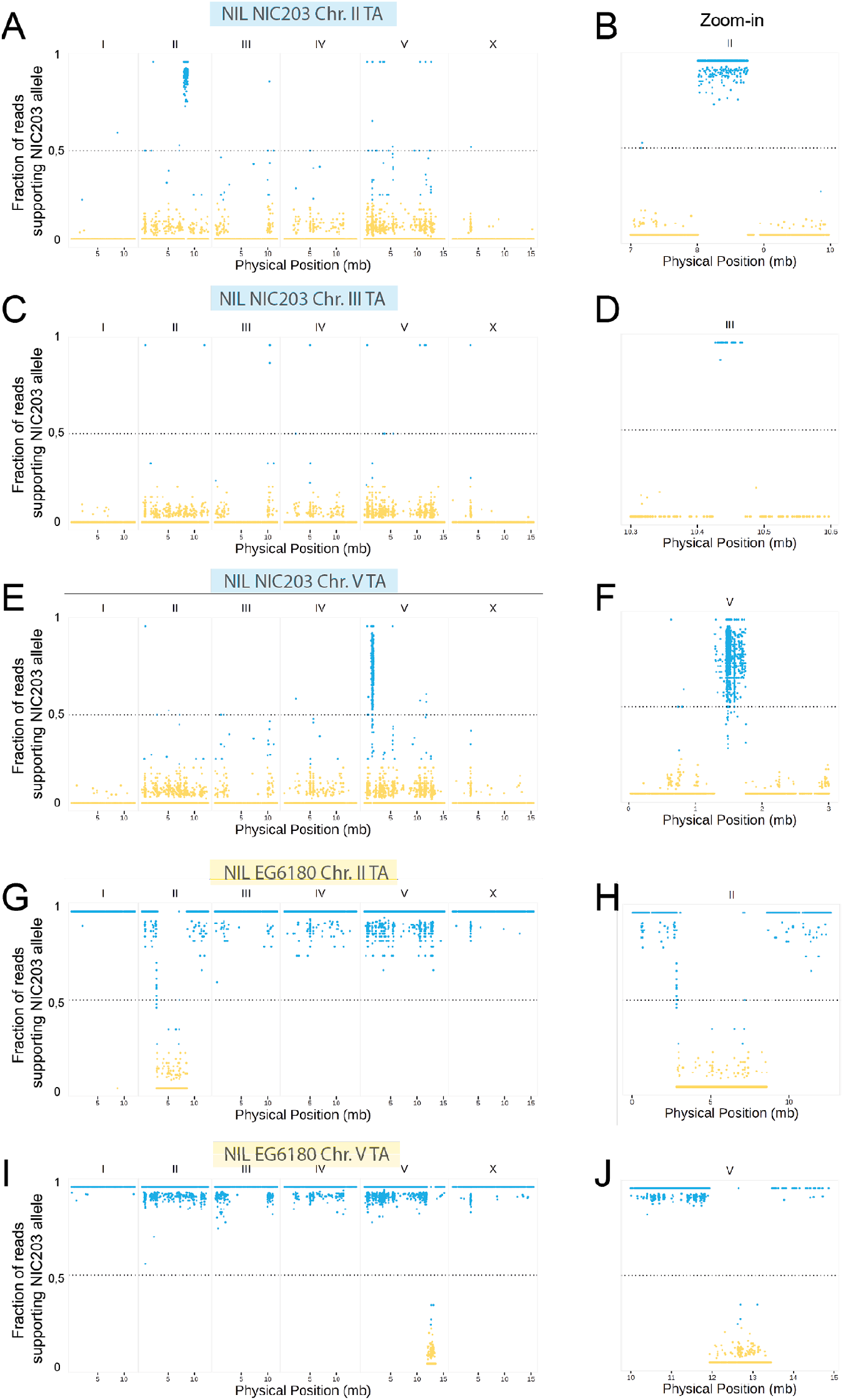
Whole-genome sequencing of NILs confirms introgressions. We performed Illumina short-read whole-genome sequencing to confirm NILs carrying only one TA element per introgressed region. Reads were aligned to the NIC203 assembly genome (**A, C, E, G, I**). Highlighted zoom-in regions of introgressions for each NIL (**B, D, F, H, J**). The NIC203 Chr. III NIL carries a very small introgression, only 40 Kb, which is not obvious in (C) but can be easily appreciated in (D).

**Figure S6.**
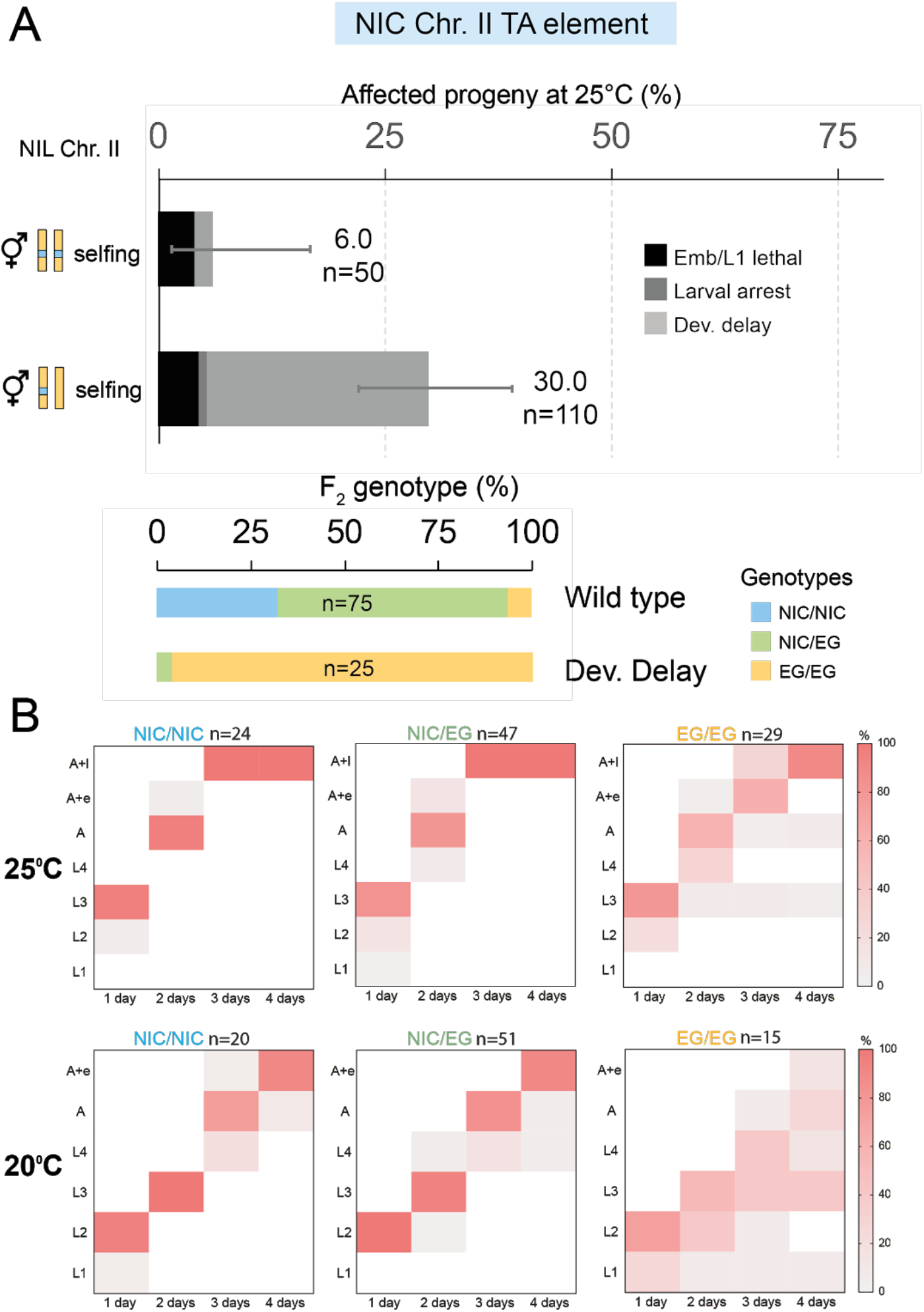
NIC203 Chr. II TA has a higher expressivity at lower temperatures. **(A)** NIC203 Chr. II TA is also active at 25℃. The parental NIL only shows background levels of developmental defects (6%, n=50), whereas ~25 % of the F_2_ progeny from a NIC Chr. II NIL x EG6180 cross is developmentally delayed (30%, n=110). Errors bars indicate 95% confidence intervals calculated with the Agresti-Coull method (*40*). The vast majority of delayed F_2_ progeny are homozygous EG/EG non-carriers. **(B)** Developmental timing of the F_2_ progeny from NIC203 Chr. II NIL x EG6180 cross for each genotype class. Single F_2_ embryos from heterozygous mothers were placed on 5cm plates seeded with OP50 at 25℃ (top) or 20℃ (top) and developmental stages were scored every 24 hours (L1-L4=larval stages, A=Adults, A+e=Adults that laid eggs, A+l=Adults and larvae progeny) n is the number of genotyped worms per class. At 20℃, the developmental delay homozygous EG/EG F_2_ progeny has a higher expressivity.

**Figure S7.**
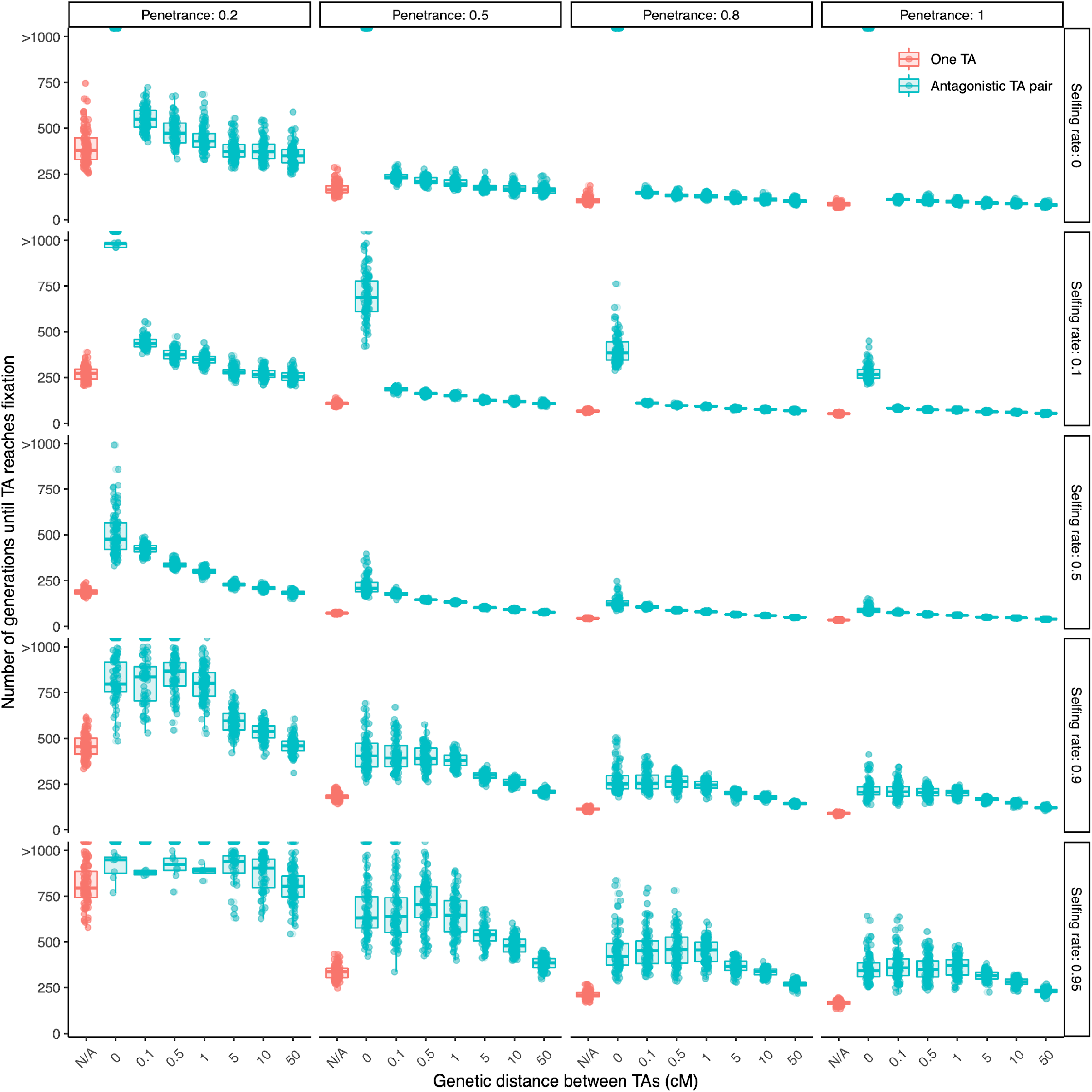
Simulations of single and antagonizing TAs across a wide range of parameters. We simulated a single TA (red) or antagonizing TAs (green) driving in a population starting at 50% allele frequency. We explored the effect of the following variables: selfing rate, penetrance of the TA, and genetic distance (for TA pair only). Across a wide range of parameters, antagonizing TA pair take longer to reach fixation in the population.

**Figure S8.**
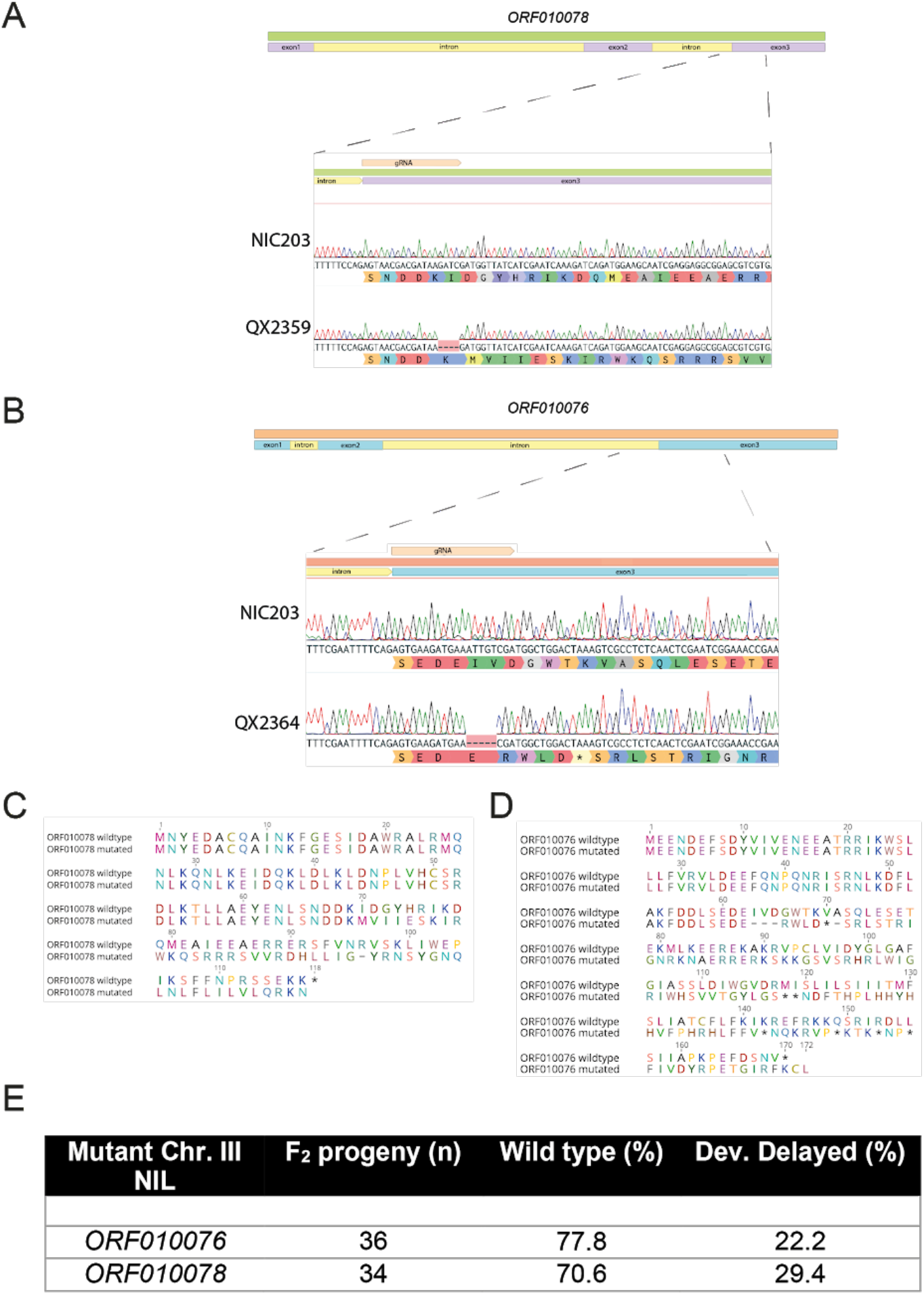
Mutations in *ORF010078* and *ORF010076* do not abolish NIC203 Chr. III TA associated developmental delay. **(A)** The *ORF010078* mutant allele generates a frameshift in the third exon, which affects the protein starting in the amino acid position 70 (length of WT is 117 aa) **(B)** The *grow-1* mutant allele introduces a frameshift and premature stop codon in the third exon, resulting in a 66 aa peptide (wild type is 169 aa). **(C)** Protein alignment of WT and mutant ORF010078. **(D)** Protein alignment of WT and mutant ORF010076. **(E)** We crossed NIC Chr. III NILs that carried mutations in either ORF010078 or ORF010076 to EG6180 males and scored their F_2_ progeny. In both cases we observed ~25% of developmentally delayed progeny, indicating that none of these two genes is the underlying toxin.

**Figure S9.**
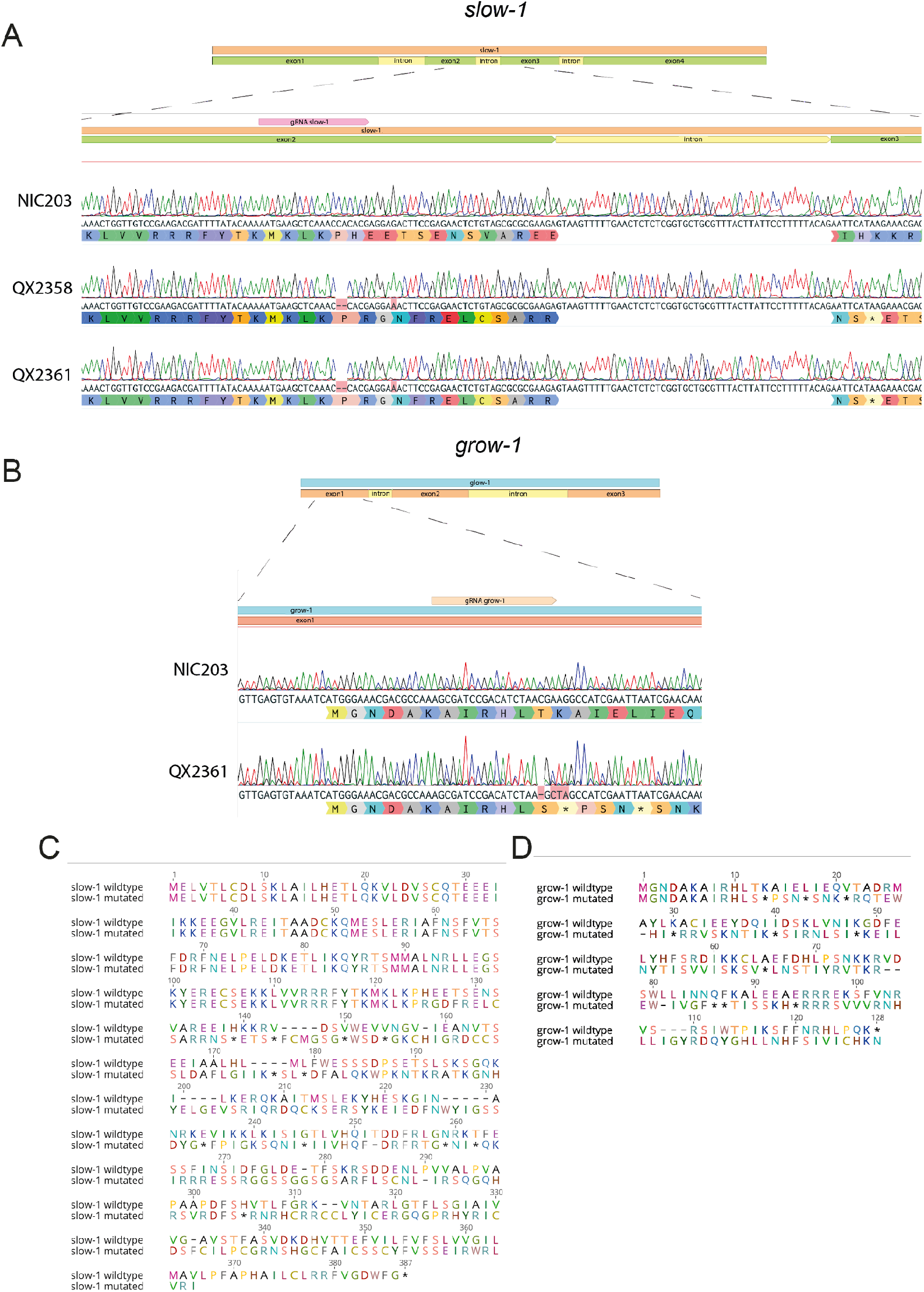
Mutant alleles of *slow-1* and *grow-1* generated using CRISPR/Cas9 gene editing. **(A)** The *slow-1* mutant allele generates a frameshift in the second exon and premature stop codon, resulting in a 129 aa peptide (wild type is 387 aa). **(B)** The *grow-1* mutant allele introduces a premature stop codon in the first exon, resulting in a 12 aa peptide (wild type is 128 aa). **(C)** Protein alignment of WT and mutant SLOW-1. **(D)** Protein alignment of WT and mutant GROW-1.

**Figure S10.**
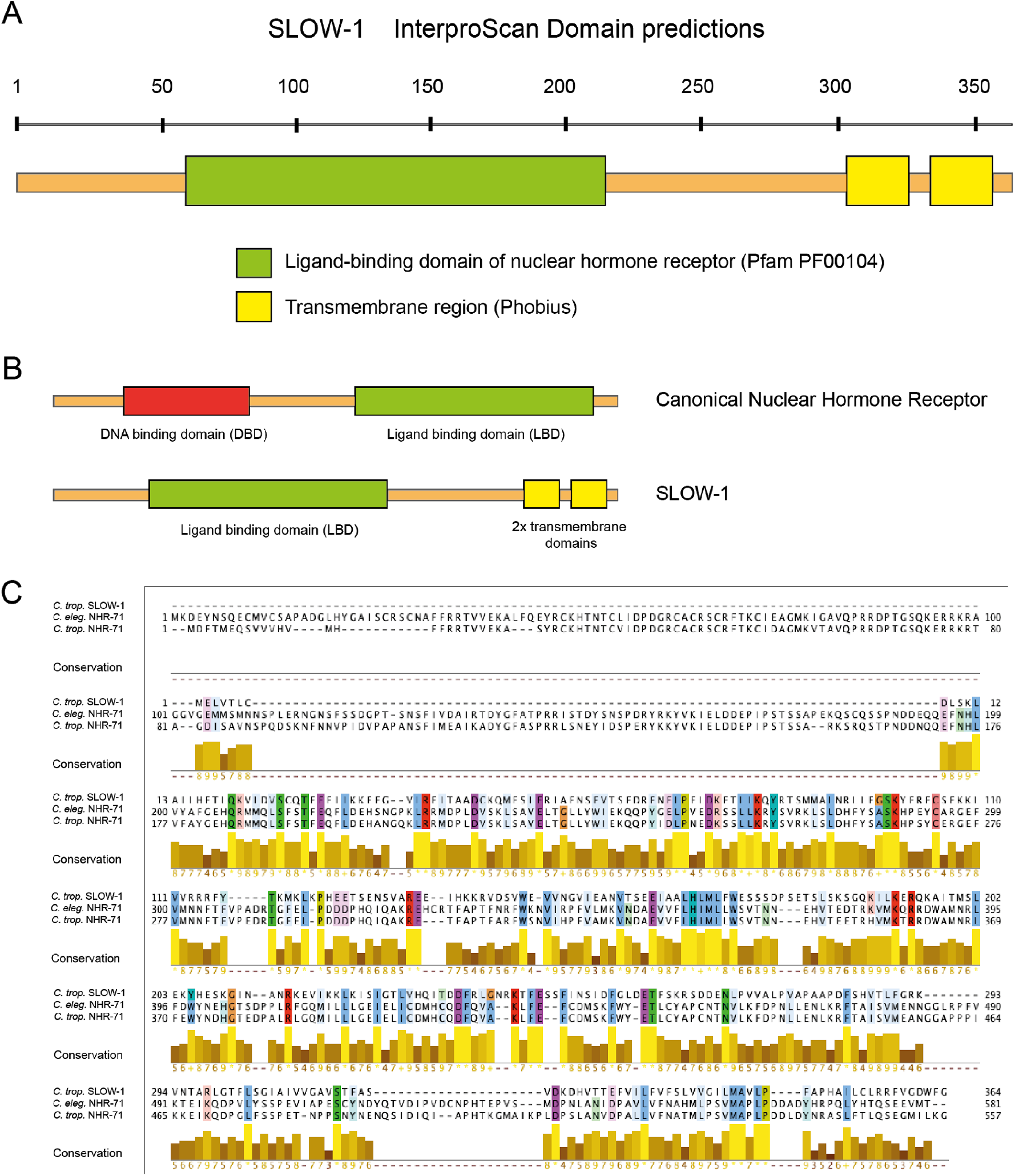
SLOW-1 is homologous to nuclear hormone receptors. **(A)** Proteins domain predicted by InterproScan **(B)** Comparison of domains found on a canonical nuclear hormone receptor (NHR) and SLOW-1. SLOW-1 is missing a DNA binding domain but contains two predicted transmembrane domains, which are not found in any other NHR to our knowledge **(C)** tblastn search of other proteins with homology to SLOW-1 in NIC203 identified NHR-71, a conserved nuclear hormone receptor, as the best blast hit (E-value 1.3). The lack of a close NHR paralog in *C. tropicalis* and other nematodes suggests that SLOW-1 is evolving under strong selection. Protein alignment of *C. tropicalis* SLOW-1, *C. elegans* NHR-71, and *C. tropicalis* NHR-71. SLOW-1 aligns to the C-terminal region of NHR-71, which contains the Ligand Binding Domain. Color code is Clustalx and highlighted residues have >75% conservation. Alignment plot generated in Jalview.

**Figure S11.**
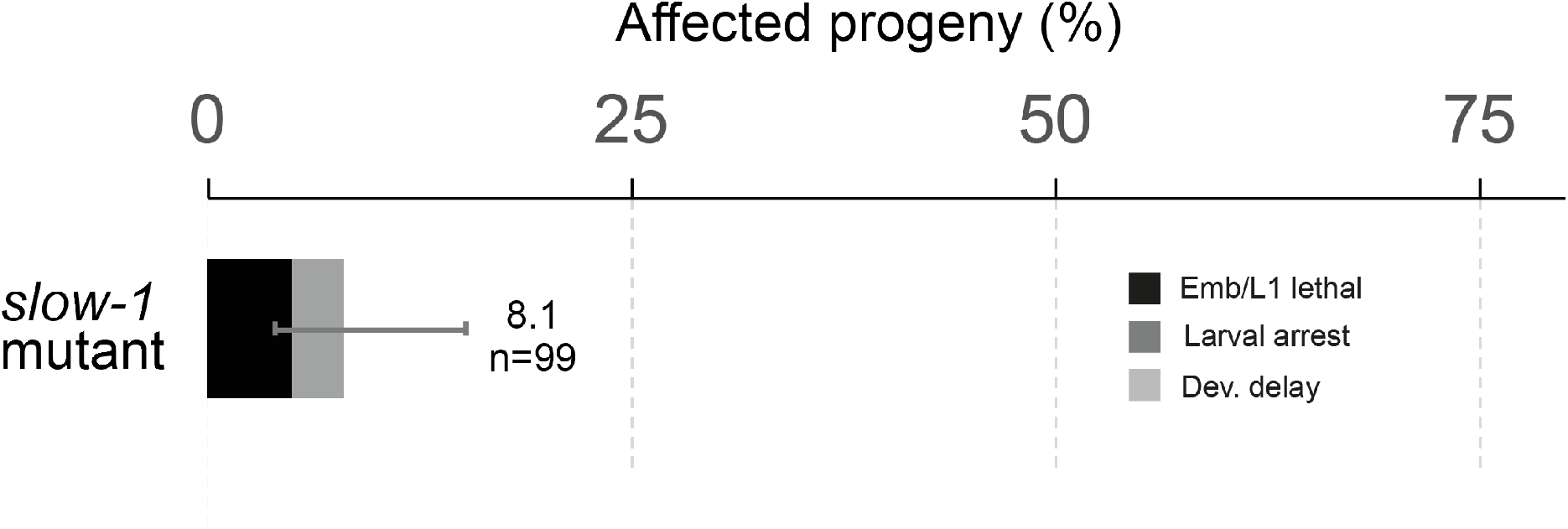
Homozygous *slow-1* mutants do not show significant developmental defects. The NIC Chr. III NIL line carrying a homozygous mutation in *slow-1* shows only background levels of defects (5% embryonic lethal and 3% larval arrest) and no developmental delay, suggesting that *slow-1* is dispensable for the development of *C. tropicalis* (n=99). Errors bars indicate 95% confidence intervals calculated with the Agresti-Coull method (*40*).

